# Pre-mRNA fate decision safeguards the fidelity of the inflammatory response

**DOI:** 10.1101/2023.11.30.569392

**Authors:** Annika Bestehorn, Jeanne Fesselet, Sebastian Didusch, Christina Zeiler, Kevin Doppelmayer, Martina Borroni, Anita LeHeron, Vera Pfanzagl, WeiQiang Chen, Sara Scinicariello, Manuela Baccarini, Gijs A Versteeg, Markus Hartl, Pavel Kovarik

## Abstract

The fidelity of immune responses depends on a timely controlled and selective mRNA degradation that is largely driven by RNA-binding proteins (RBPs)[1, 2]. It remains unclear whether the selection of an individual mRNA molecule for degradation is governed by stochastic or directed processes. Here, we show that tristetraprolin (TTP, also known as ZFP36), an essential anti-inflammatory RBP[3], destabilizes target mRNAs via a hierarchical molecular assembly. The formation of the assembly strictly relies on the interaction of TTP with RNA. The TTP homolog ZFP36L1 exhibits similar requirements, indicating a broader relevance of this regulatory program. Unexpectedly, the assembly of the cytoplasmic mRNA-destabilization complex is licensed in the nucleus by TTP binding to pre-mRNA, while mature cytoplasmic mRNA does not constitute a *de novo* TTP target. Hence, the fate of an inflammation-induced mRNA is decided concomitantly with its synthesis. This decision mechanism prevents the translation of superfluous and potentially harmful inflammation mediators, ensuring an efficient cessation of the immune response, irrespective of transcriptional activity.

## INTRODUCTION

Functional immune responses are dependent on rapid and precisely controlled adjustments in the transcriptome and proteome to ensure protection against pathogens while minimizing damage to the host. A key safeguarding mechanism against an aggravated immune response is the orchestrated and selective destabilization of mRNAs coding for inflammatory mediators [2]. The major drivers of mRNA destabilization are cis-acting RNA-binding proteins (RBPs), which act mostly as adaptors for the RNA degradation machinery, notably the CCR4-NOT deadenylation and the DCP decapping complexes [1, 4–6]. The selectivity in mRNA destabilization is defined by the interaction of an RBP with a sequence or structure in the target RNA molecule. Although this basic concept of RBP-driven mRNA destabilization is established, the hierarchy of the process is not understood. Consequently, the elementary question of whether the binding of a given RBP to an individual mRNA molecule is a stochastic event or whether it occurs in a defined phase of the mRNA life cycle, remains to be answered.

To determine the molecular hierarchy of mRNA decay orchestrating the resolution of inflammation we employed TTP, which is an mRNA-destabilizing RBP with essential functions in the control of inflammatory gene expression [7]. TTP deletion in mice causes multi-organ inflammation including arthritis, dermatitis, and cachexia, cumulatively known as the TTP deficiency syndrome [3]. TTP binds to adenine-uridine-rich elements (AREs) in 3’ untranslated regions (UTRs) via its tandem zinc fingers (TZF) and interacts with both CCR4-NOT and DCP complexes, thereby causing destabilization of target mRNAs [8–13]. TTP targets are enriched in, but not limited to, mRNAs coding for cytokines and chemokines [13–15]. The TTP deficiency phenotype is causally associated with increased stability and expression of *Tnf*, *Il23*, *Il1a*, *Ccl3*, and possibly other cytokine or chemokine mRNAs [3, 16–18]. External stress cues that activate the p38 MAP kinase induce MK2-mediated TTP phosphorylation, thereby increasing TTP protein stability but inhibiting its mRNA-destabilization activity [19, 20]. The organization and order of individual steps of TTP-dependent RNA decay within the mRNA life cycle remain unknown. Filling this knowledge gap is essential for the understanding of non-resolving inflammation, which is often causative of autoimmunity, cancer or neurodegenerative disorders.

## RESULTS

### RNA binding enables TTP to associate with mRNA degradation complexes and prevents continuous TTP accumulation in the cell nucleus

The cardinal feature of TTP is its ability to orchestrate the timing, extent and selectivity of mRNA decay required for efficient but not exaggerated immune responses. To understand the hierarchy of this process, we first determined molecular assemblies representing distinct steps of TTP-mediated mRNA decay using APEX2-based proximity-dependent labeling. APEX2 is an engineered ascorbate peroxidase with an improved labeling activity [21]. We fused the murine wild-type (WT) TTP (GenBank NP_035886) or an RNA binding-deficient tandem zinc finger TTP mutant (TZF^mut^) N-terminally to APEX2 to obtain WT-APX and TZF^mut^-APX constructs allowing inducible expression with the tetracycline derivative doxycycline (**Fig. 1a**). TZF^mut^ was constructed by introducing the mutations C116R and C139R, which inactivate the zinc-coordinating properties of each of the two zinc fingers and abolish RNA binding, as reported previously [22]. A GFP-APX control was generated to determine the background labeling. The constructs were introduced into HEK293 cells using lentiviral transduction, and their doxycycline-inducible expression under steady state and stress conditions was validated (**Fig. 1b**). Stress conditions were imposed using anisomycin which, similar to other stress cues, increases phosphorylation of TTP at S52 and S178 (mouse coordinates) [23–25]. Anisomycin treatment resulted in a slower mobility of a proportion of TTP fusion proteins, indicative of increased phosphorylation [24] (**Fig. 1b**).

**Figure 1.**
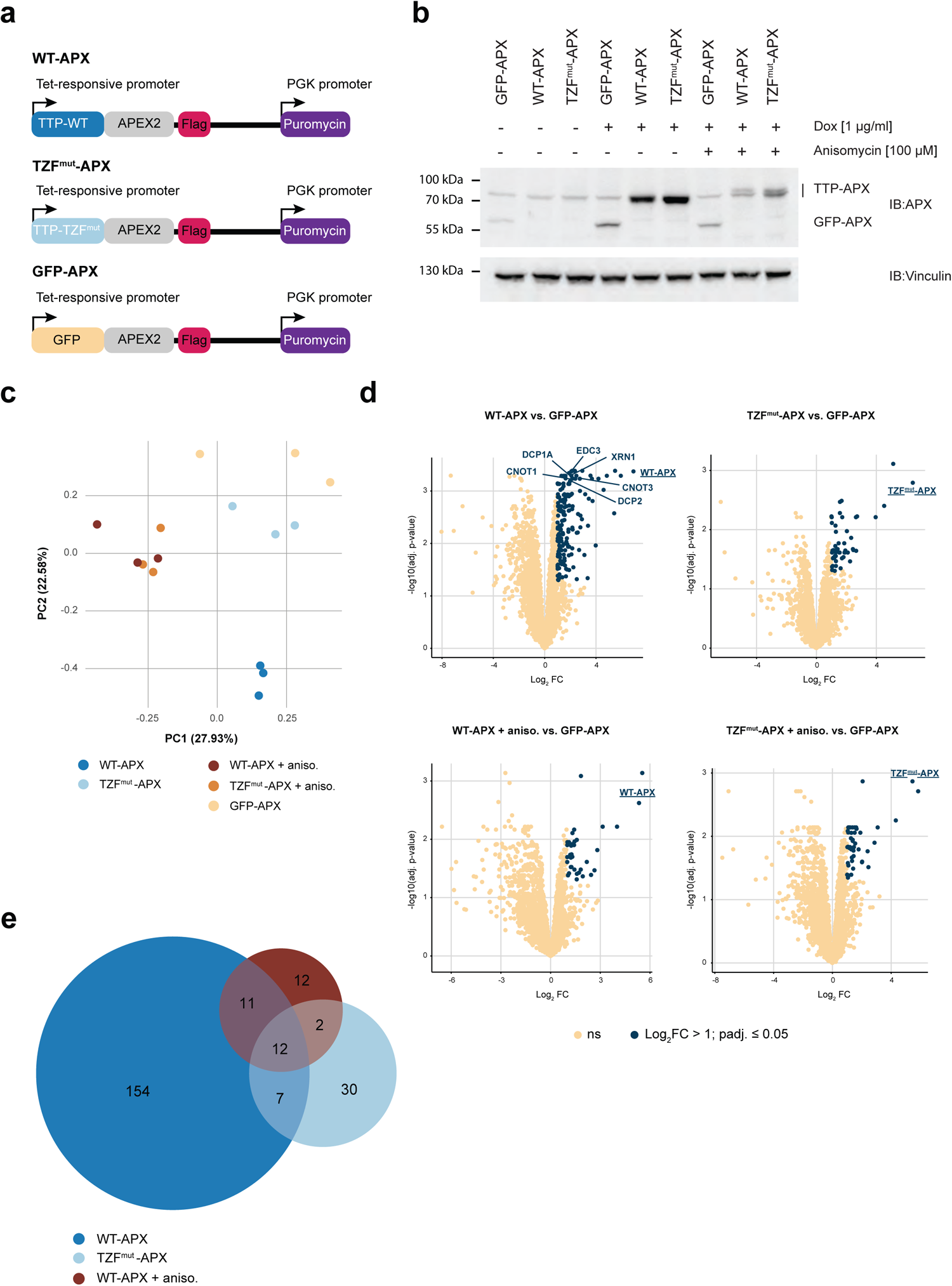
Effects of RNA binding and external stress cues on TTP proximal interactome. **a.** Schematic representation of constructs used in proximity labeling experiments. Wild-type TTP, RNA binding-deficient tandem zinc finger TTP mutant (TZF^mut^) and GFP fused N-terminally to APEX2 downstream of a tetracycline-responsive promoter yielding WT-APX, TZF^mut^-APX and GFP-APX constructs. All constructs contained a C-terminal Flag tag and a puromycin resistance cassette. **b.** Inducible expression of WT-APX, TZF^mut^-APX and GFP-APX. Expression of WT-APX, TZF^mut^-APX and GFP-APX in HEK293 cells was induced by doxycycline for 3.5 h. Anisomycin (1.5 h) was used as stress stimulus, as indicated. Whole cell lysates were analyzed by Western blotting using APEX antibodies (immunoblot, IB: APX); vinculin (IB: Vinculin) served as a loading control. Positions of TTP-APX fusion proteins (WT-APX and TZF^mut^-APX) and GFP-APX are indicated. Note that anisomycin treatment resulted in the appearance of slower migrating TTP-APX fusions, corresponding to more highly phosphorylated TTP fusion proteins. **c - e.** Analysis of APX-based proximal TTP interactome in HEK293 cells by LC-MS/MS. Expression of WT-APX, TZF^mut^-APX and GFP-APX was induced by doxycycline as in (b). Anisomycin treatment (+aniso) was employed as in (b). Following labeling with biotin-phenol and H_2_O_2_, biotinylated proteins were enriched by streptavidin pull-down and analyzed by LC-MS/MS. (**c**) Principal component analysis (PCA) of protein intensities visualized clustering of biological triplicates and effects of RNA binding (TZF^mut^-APX) and stress stimulation (+aniso) on the proximal proteome. (**d**) Differential enrichment analysis for the comparisons of interactomes WT-APX vs. GFP-APX, TZF^mut^-APX vs. GFP-APX, WT-APX+aniso vs. GFP-APX, and TZF^mut^-APX+aniso vs. GFP-APX. Significantly enriched proteins are depicted in blue (log_2_ fold change (FC) > 1, padj. ≤ 0.05, n = 3). WT-APX, TZF^mut^-APX and GFP-APX fusion constructs are underlined. Known TTP interactors DCP1A, DCP2, EDC3, CNOT1, CNOT3 and XRN1 are annotated; bold blue indicates significant enrichment. Note that these interactors were not significantly enriched in the TZF^mut^-APX, WT-APX+aniso and TZF^mut^-APX+aniso interactomes (**Supplementary Table 2**). (**e**) Overlap of proteins enriched in WT-APX, TZF^mut^-APX, and WT-APX+aniso interactomes over GPF-APX background labeling control (Log2FC > 1, padj. ≤ 0.05, n = 3).

Proximity labeling was performed by the addition of biotin-phenol for 30 min and subsequent treatment with H_2_O_2_ for 1 min. Successful labeling in cells expressing WT-APX, TZF^mut^-APX or GFP-APX was verified by the detection of increased levels of biotinylated proteins in whole-cell-extract samples and streptavidin pull-down samples from H_2_O_2_-treated cells, as compared to samples from untreated cells (**Extended Data Fig. 1a**). The presence of biotinylated proteins in cells not treated with H_2_O_2_ resulted from endogenous biotin-dependent carboxylases and endogenous biotinylation. Three replicates of labeled pull-down samples for each of the three constructs (i.e. WT-APX, TZF^mut^-APX or GFP-WT) were generated. In addition, samples from anisomycin-treated WT-APX- or TZF^mut^-APX-expressing cells (WT-APX+aniso and TZF^mut^-APX+aniso) were prepared to estimate the effect of TTP phosphorylation on the proximal interactome. The streptavidin pull-down samples were subject to LC-MS/MS analysis to identify and quantitate biotinylated proteins (**Supplementary Table 1**). The numbers of identified proteins were similar in all sample replicates (**Extended Data Fig. 1b**). Principal component analysis (PCA) of protein intensities revealed distinct clusters of the corresponding replicates (**Fig. 1c**). The WT-APX cluster was remarkably well separated from the TZF^mut^-APX cluster, indicating that binding of TTP to RNA was a major determinant of the TTP interactome. Anisomycin treatment changed the position of both WT-APX and TZF^mut^-APX clusters. Surprisingly, anisomycin caused both WT-APX and TZF^mut^-APX samples to cluster in the same area (**Fig. 1c**), suggesting that stress-induced phosphorylation overruled the differences between the interactomes of wild-type and RNA binding-deficient TTP.

To identify proteins proximally interacting with TTP, we determined the enrichment over the GFP-APX sample for each TTP construct and condition using the amica proteomics data analysis platform [26]. WT-APX displayed 184 enriched proteins, while the number of enriched proteins was substantially lower in TZF^mut^-APX (51 proteins), WT-APX+aniso (37 proteins) and TZF^mut^-APX+aniso (50 proteins) samples (**Supplementary Table 2, Fig. 1d**). The WT-APX interactome comprised the XRN1 exonuclease as well as the deadenylation and decapping components CNOT1, CNOT3, DCP1, DCP2, and EDC3 (**Fig. 1d**) that were previously reported to bind to TTP [9–11, 27, 28]. Strikingly, these proteins were missing in the TZF^mut^-APX sample suggesting that TTP interacts with the deadenylation and decapping enzymes only following its binding to RNA. The preferential association of CNOT1, DCP1a and EDC3 with WT-APX as compared to TZF^mut^-APX was verified by Western blotting (**Extended Data Fig. 1c**).

Consistent with the reported inhibitory effect of TTP phosphorylation on its binding to the deadenylase complex [19], anisomycin treatment abolished WT-APX interactions with the deadenylase subunits (e.g. CNOT1 and CNOT3) (**Fig. 1d**). Moreover, TTP interactions with the decapping complex were also lost upon anisomycin treatment (**Fig. 1d**).

The critical impact of RNA binding and anisomycin treatment on the TTP interactome was further illustrated by the limited overlap of proteins enriched in WT-APX, TZF^mut^-APX and WT-APX+aniso samples (**Fig. 1e**). Anisomycin changed the divergent WT-APX and TZF^mut^-APX interactomes into highly similar ones, with WT-APX+aniso and TZF^mut^-APX+aniso samples sharing 31 proteins (out 37 and 50, respectively) indicating the acquisition of a uniform proximal environment by RNA-bound and RNA-unbound TTP upon stress (**Extended Data Fig. 1d**).

Top-ranked Gene Ontology (GO) terms significantly overrepresented in the WT-APX interactome, i.e. in proteins enriched in WT-APX over GFP-APX profiles, comprised terms consistent with the established TTP function: GO cellular compartment (CC) included “cytoplasmic RNP granule” and “P-body” terms, GO biological process (BP) included “posttranscriptional regulation of gene expression”, “mRNA catabolic process”, and “regulation of RNA stability” terms, and GO molecular function (MF) included “RNA binding” and “mRNA 3’ UTR binding” terms (**Fig. 2a**). In contrast, no such GO terms were found in the TZF^mut^-APX interactome which instead comprised the terms “nucleus” and “nucleoplasm” in the CC category, suggesting a nuclear localization; the category BP contained only one term and no terms were found in the MF category (**Fig. 2a**). Only few and inconsistent GO terms were overrepresented in the anisomycin-dependent TTP interactomes (WT-APX+aniso and TZF^mut^-APX+aniso) (**Extended Data Fig. 2a**). Overall, the GO term analysis was conclusive for the WT-APX and TZF^mut^-APX interactomes: the terms associated with WT-APX allowed inference of the function of TTP in mRNA destabilization while, surprisingly, the terms associated with TZF^mut^-APX suggested that the RNA binding-deficient TTP localizes in the nucleus (**Fig. 2b**). These conclusions were underpinned by the analysis of the terms “P-body” and “Nucleus” at the gene level showing that the TZF^mut^-APX interactome was depleted in almost all “P-body” genes while it was enriched in many “Nucleus” genes (**Extended Data Fig. 2b**). Importantly, several “Nucleus” genes were enriched also in the WT-APX interactome (e.g. ATAD5, NFATC1, and MAGEB2) indicating that the WT-APX protein was not excluded from the nucleus (**Extended Data Fig. 2b**), consistent with the reported predominantly cytoplasmic localization and nucleo-cytoplasmic shuttling of TTP [24, 29]. The different cellular localization of WT-APX and TZF^mut^-APX was further supported by higher counts of the known cytoplasmic TTP interactor 14-3-3 protein [24] in WT-APX as compared to TZF^mut^-APX interactomes (YWHAQ protein log_2_FC = 0.52, padj = 0.005; **Supplementary Table 1**).

**Figure 2.**
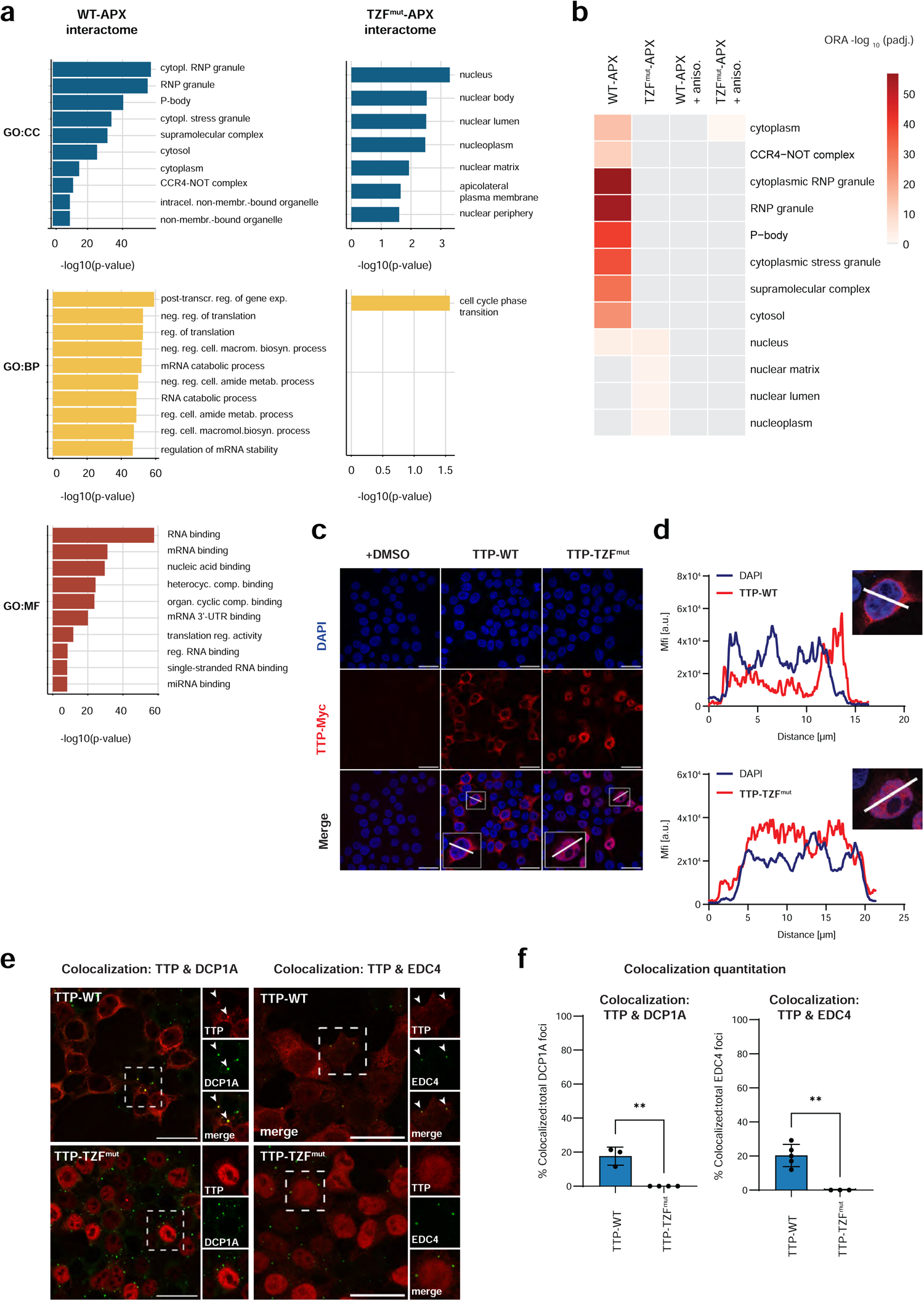
Effects of RNA binding on association of TTP with mRNA degradation complexes and TTP subcellular localization. **a.** Overrepresentation of gene ontology (GO) terms in WT-APX and TFZ^mut^-APX interactomes. Interactomes were defined as proteins significantly enriched over GFP-APX background labeling control (log_2_FC ≥ 1 and padj. < 0.05, n = 3) in LC-MS/MS data. Bar plots depict up to top 10 most significantly overrepresented terms in the GO categories Cellular compartment (CC), Biological process (BP) and Molecular function (MF). Note that the TZF^mut^-APX interactome did not contain any enriched terms in the MF category and is therefore not shown. **b.** Comparison of top eight GO CC terms in the WT-APX interactome and top four GO CC terms in the TZF^mut^-APX interactome, across all four analyzed interactomes. The CC terms were selected from terms identified in (a); padj. values of overrepresentation analysis (ORA) shown in color code; grey color: not significant. **c, d.** Immunofluorescence microscopy of C-terminally Myc-tagged TTP-WT and TTP-TZF^mut^ in HEK293 cells. Expression of TTP constructs was induced by doxycycline (1 µg/ml, 3.5 h); control cells were treated with vehicle (DMSO). Constructs were detected with a Myc antibody; DAPI visualized the nucleus. (**c**) Immunofluorescence images with profile lines drawn through representative cells (enlarged 2-fold in insets) for quantitation of mean fluorescence intensity (MFI) in (d). Scale bar = 25 µm. (**d**) Quantitation of MFI along a profile line drawn through representative cells in (c). **e, f.** Colocalization of TTP-WT or TTP-TZF^mut^ with DCP1A and EDC4 foci. TTP-WT and TTP-TZF^mut^ were expressed as in (c). (**e**) Immunofluorescence images showing TTP-WT or TTP-TZF^mut^ and endogenous DCP1A and EDC4. Large images depict TTP constructs visualized with a Myc antibody (red); highlighted cells are shown in insets together with DCP1A staining or EDC4 staining (both green) in individual and merged channels. Arrowheads point to foci of DCP1A or EDC4 colocalizing with TTP signal. Scale bar = 25 µm. (**f**) Quantitation of colocalization of TTP-WT or TTP-TZF^mut^ signals with DCP1A or EDC4 foci (Fiji plugin Comdet, see Methods). Student’s t test. **padj. < 0.01, n = 60 −113 cells

The results of proximity labeling experiments were validated by immunofluorescence microscopy using HEK293 cells expressing myc-tagged TTP-WT or TTP-TZF^mut^ proteins. Consistent with the GO term analysis, TTP-TZF^mut^ was confined to the nucleus and TTP-WT localized mostly but not exclusively in the cytoplasm (**Fig. 2c, d**). The difference in the localization pattern was significant (**Extended Data Fig. 2c**). The proximity labeling results were further validated by double immunofluorescence staining showing colocalization of TTP-WT with DCP1a and EDC4 foci (**Fig. 2e, f**), which represent cytoplasmic RNP granules possessing decapping activity [30]. No colocalization of TTP-TZF^mut^ with the DCP1a and EDC4 foci was detected, in agreement with the proximal interactome (**Fig. 2e, f**).

Cumulatively, these results reveal that RNA binding has a regulatory in addition to the canonical, i.e. RNA binding, function in TTP: RNA binding is required for the maintenance of the cytoplasmic TTP pool and for the association of TTP with RNA degradation complexes.

### The regulatory function of RNA binding is conserved within the TTP protein family and is effective also in immune cells

TTP is member of a conserved family of TZF proteins which includes ZFP36L1 and ZFP36L2 [31]. The sequence similarity within this protein family is the highest in the TZF domain (approximately 70% identity) (**Extended Data Fig. 3a**). To test whether the role of RNA binding in the interaction with RNA processing enzymes and in the subcellular localization is conserved in the TTP protein family, we determined the proximal interactomes and subcellular localization of ZFP36L1 and its RNA binding-deficient derivative using the same strategy as for TTP. We generated the APX fusion proteins L1 WT-APX and L1 TZF^mut^-APX, inducibly expressed them in HEK293 cells and confirmed their labeling activity (**Extended Data Fig. 3b, c**). Following the treatment of L1 WT-APX-, L1 TZF^mut^-APX- and GFP-APX-expressing cells with biotin-phenol and H_2_O_2_, peptide intensities in streptavidin pull-downs were determined by LC-MS/MS (**Supplementary Table 3**). The replicates of each L1 WT-APX, L1 TZF^mut^-APX and GFP-APX clustered in separated areas in PCA (**Extended Data Fig. 3d**), similar to TTP samples. Also comparable to the TTP experiments, analysis of proteins enriched in L1 WT-APX or L1 TZF^mut^-APX samples over GFP-APX revealed a higher number of interaction partners in the L1 WT-APX interactome than in the L1 TZF^mut^-APX sample (283 versus 38) (**Supplementary Table 4**, **Fig. 3a**). Importantly, proteins found in the L1 WT-APX interactome comprised subunits of the deadenylation and decapping complexes (e.g. CNOT3, EDC3, and EDC4) while these proteins were missing in the L1 TZF^mut^-APX interactome (**Fig. 3a**).

**Figure 3.**
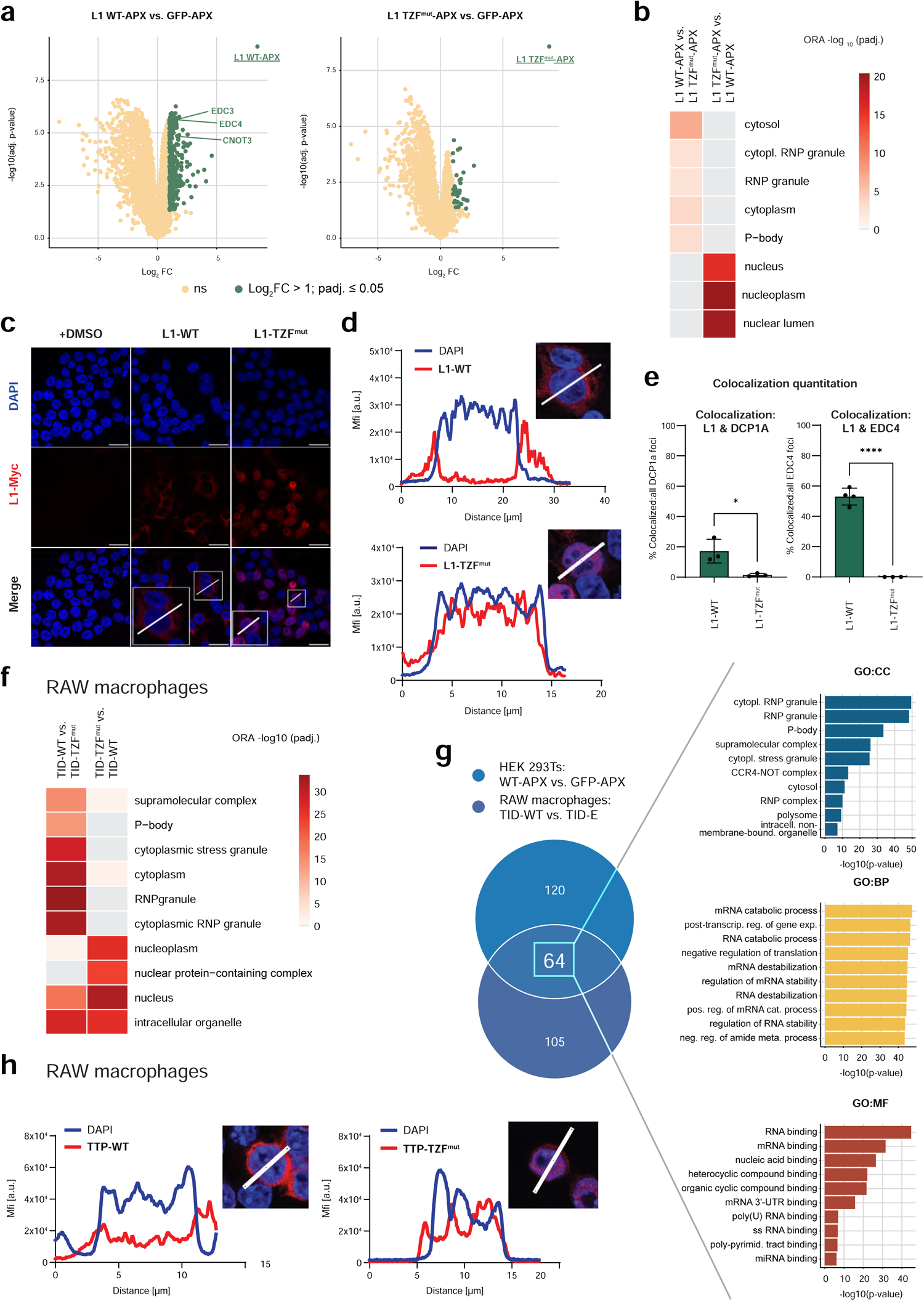
Role of RNA binding in the interactome and subcellular localization of ZFP36L1 in HEK293 cells and of TTP in macrophages. **a.** Analysis of APX-based proximal ZFP36L1 interactome in HEK293 cells by LC-MS/MS. Expression of ZFP36L1 WT-APX (L1 WT-APX), ZFP36L1 TZF^mut^-APX (L1 TZF^mut^-APX) and GFP-APX and analysis of biotinylated proteins by LC-MS/MS was carried out as described for TTP constructs in Fig. 2. Differential enrichment analysis defining the L1 WT (L1 WT-APX vs. GFP-APX) and L1 TZF^mut^-APX (L1 TZF^mut^-APX vs. GFP-APX) interactomes. Significantly enriched proteins are depicted in blue (log_2_FC > 1, padj ≤ 0.05, n = 3). L1 WT-APX and L1 TZF^mut^-APX fusion constructs are underlined. The TTP interactors EDC3, EDC4 and CNOT3 are annotated; bold blue indicates significant enrichment. Note that these interactors were not significantly enriched in the L1 TZF^mut^-APX interactome (**Supplementary Table 3**). **b.** GO term (CC category) overrepresentation in proteins enriched in L1 WT-APX interactome (L1 WT-APX vs. L1 TZF^mut^-APX) and in L1 TZF^mut^-APX interactome (L1 WT-APX vs. L1 TZF^mut^-APX). ORA padj. values shown in color code; grey color: not significant. **c, d.** Immunofluorescence microscopy of C-terminally Myc-tagged L1-WT and L1-TZF^mut^ in HEK293 cells. Expression of L1 constructs was induced by doxycycline (1 µg/µl, 3.5 h); control cells were treated with vehicle (DMSO). Constructs were detected with a Myc antibody; DAPI visualized the nucleus. (**c**) Immunofluorescence images with profile lines drawn through representative cells (enlarged 2-fold in insets) for quantitation of mean fluorescence intensity (MFI) in (d). Scale bar = 25 µm. (**d**) MFI profile of Myc and DAPI signals along a line drawn through representative cells depicted in (c). The MFI profiles of Myc and DAPI signals were similar in L1-TZF^mut^ implying nuclear localization of the protein. **e.** Colocalization of L1-WT or L1-TZF^mut^ with DCP1A and EDC4 foci. L1-WT and L1-TZF^mut^ were expressed as in (c). L1 constructs were detected using a Myc antibody, endogenous DCP1A and EDC4 were detected using DCP1A and EDC4 antibodies, and colocalization was quantitated (Fiji plugin Comdet). Student’s t test. **padj. < 0.01, n = 44-132 cells. **f.** GO term overrepresentation in proximal TurboID (TID)-based interactomes of TTP WT (TID-WT) and TTP TZF^mut^ (TID-TZF^mut^) in RAW macrophages deficient in endogenous TTP (hereafter referred to as RAW macrophages). Expression of TID-WT and TID-TZF^mut^ in RAW macrophages was induced by doxycycline (24 h) followed by biotin labeling for 15 min. Biotinylated proteins were analyzed by LC-MS/MS. GO terms overrepresented in TID-WT and TID-TZF^mut^ interactomes (defined by comparisons TID-WT vs. TID-TZF^mut^ or TID-TZF^mut^ vs. TID-WT, respectively) were determined using ORA. Top eight GO CC terms in TID-WT and top four GO CC terms in TID-TZF^mut^ are shown. ORA padj. values shown in color code; grey color: not significant. **g.** Overlap of TTP interactomes in HEK293 cells (WT-APX vs. GFP-APX) and RAW macrophages (TID-WT vs. TID-E) shown in Venn diagram. Proteins in overlap (64 interactors) were subjected to GO term ORA (log_2_ FC > 1 and padj. ≤ 0.05). **h.** Subcellular localization of TTP-WT and TTP-TZF^mut^ in RAW macrophages. TTP-WT and TTP-TZF^mut^ expression in RAW macrophages was induced with doxycycline (0.5 µg/ml, 24 h), and localization of TTP constructs was determined by immunofluorescence microscopy using a Myc antibody; DAPI visualized the nucleus. Shown is MFI profile of Myc and DAPI signals along a line drawn through representative cells depicted in **Extended Data Fig. 3i**. The MFI profiles of Myc and DAPI signals were similar in TTP-TZF^mut^ implying nuclear localization of the protein.

GO terms overrepresented in proteins enriched in the L1 WT-APX interactome over the L1 TZF^mut^-APX interactome were consistent with RNA degradation (e.g. “cytoplasmic RNP granule” and “P-body”). These terms were absent from the L1 TZF^mut^-APX interactome (L1 TZF^mut^-APX versus L1 WT-APX), which comprised instead terms consistent with nuclear localization (“nuclear lumen” and “nucleus”) (**Fig. 3b**). Immunofluorescence microscopy confirmed L1-TZF^mut^ to localize in the nucleus while the L1-WT protein was predominantly cytoplasmic (**Fig. 3c and d, Extended Data Fig. 3e**). Furthermore, similar to TTP, L1-WT co-localized with DCP1a and EDC4 foci whereas L1-TZF^mut^ did not (**Fig. 3e** and **Extended Data Fig. 3f**). Thus, RNA binding enables the association of ZFP36L1 with the RNA processing complexes and prevents the protein from nuclear accumulation.

The control of inflammatory responses by TTP is most evident in immune cells such as macrophages, which abundantly express the physiological TTP targets including cytokine and chemokine mRNAs [13, 14, 32]. Therefore, to examine the regulation of TTP by RNA binding in a relevant system, we employed the RAW 264.7 murine macrophage cell line. Instead of APEX2 we used the TurboID proximity labeling system [33] due to the high level of unspecific APEX2-driven labeling we detected in these cells. We generated RAW 264.7 macrophages deficient in endogenous TTP (hereafter referred to as RAW macrophages) using CRISPR/Cas9. Subsequently, these cells were transduced with lentiviral constructs for inducible expression of TTP or the RNA binding-deficient TTP mutant C-terminally fused to TurboID (TID-WT or TID TZF^mut^) (**Extended Data Fig. 3g**). The empty TurboID construct (TID-E) was introduced to determine the background labeling. Following mass spectrometry analysis of biotinylated interactomes (**Supplementary Table 5**), PCA of protein intensities showed distinct clusters of TID-WT, TID-TZF^mut^ and TID-E replicates (**Extended Data Fig. 3h**), similar to APEX2-labeled TTP and ZFP36L1 samples in HEK293 cells. The WT-TID interactome (TID-WT versus TID-TZF^mut^ comparison, **Supplementary Table 6**) showed an enrichment with RNA degradation GO terms (e.g. “cytoplasmic RNP granule” and “P-body”), while the TID-TZF^mut^ interactome was missing these terms (**Fig. 3f**). Instead, the terms overrepresented in the TID-TZF^mut^ interactome were consistent with nuclear localization or nuclear processes. Importantly, one of these terms was also overrepresented in the WT-TID interactome, in agreement with the localization of a minor TTP pool in the nucleus reported previously [24, 29].

These results were similar to those obtained for TTP-APX constructs in HEK293 cells. The consistency of APEX2-dependent TTP interactomes in HEK293 cells and TurboID-dependent TTP interactomes in RAW macrophages were corroborated by GO term analysis of proteins found in the overlap of both approaches: the terms in all three GO categories implied an RNA-destabilizing function of TTP (**Fig. 3g**, **Supplementary Table 7**). Moreover, immunofluorescence microscopy showed TTP-TZF^mut^ to localize in the nucleus while the TTP-WT protein was mostly cytoplasmic (**Fig. 3h, Extended Data Fig. 3i, j**), thereby resembling the localization seen in HEK293 cells.

In summary, the APEX2- and TurboID-based proximity labeling approaches in HEK293 cells and RAW macrophages (**Figs. 1 - 3**) uncovered an immense impact of RNA binding on the interaction landscape of TTP. The results revealed a hierarchical molecular assembly, in which RNA binding is a prerequisite for association of TTP with the RNA degradation machinery. Moreover, RNA binding-deficient TTP was found to accumulate in the nucleus suggesting that TTP requires RNA binding for its nucleo-cytoplasmic shuttling and the acquisition of the predominantly cytoplasmic localization. We obtained similar data for ZFP36L1, indicating that the regulatory function of RNA binding is conserved in the TTP protein family.

### The primary target of TTP is pre-mRNA not mRNA

The requirement for RNA binding in TTP nucleo-cytoplasmic shuttling and association with RNA degradation enzymes prompted us to hypothesize that the initial interaction between TTP and its target RNA occurs in the nucleus and that this primary TTP-RNA complex is subsequently exported to the cytoplasm. This hypothesis is supported by previous CLIP-seq experiments carried out in macrophages and HEK293 cells, which unequivocally showed the ability of TTP to bind to AREs in introns, in addition to those in 3’ UTRs [13, 14, 34]. We thus reasoned that the principal TTP target is pre-mRNA rather than mature mRNA.

To test this hypothesis, we determined the ratios of pre-mRNA versus mRNA in the TTP-bound RNA pool and compared them to the ratios in the total cellular RNA. If the prediction that TTP primarily targets pre-mRNA was correct, then the pre-mRNA to mRNA ratios should be higher in the TTP-bound RNA pool than in the total cellular RNA (**Fig. 4a**). To isolate TTP-bound full-length RNA molecules, we modified the thiouridine (4sU)-facilitated CLIP-seq approach used by us previously [13]: we omitted the limited RNA hydrolysis step after the pull-down such that the TTP-bound RNA molecules were obtained and sequenced in their entire length. We expressed myc-tagged TTP in RAW macrophages devoid of endogenous TTP, and induced the expression of TTP targets with LPS for 6 h, in the presence of 4sU in the last hour of LPS stimulation. Following crosslinking with 365 nm UV, TTP-bound RNA was isolated using pull-down for myc-tagged TTP and subjected to total RNA-seq, together with the total cellular RNA (input samples). The integrity of input and pull-down samples was confirmed by read coverage plots for *Tnf* and *Cxcl2* (**Fig. 4b**) as representatives of TTP targets with well characterized TTP binding peaks mapped in a transcriptome-wide study in macrophages (https://ttp-atlas.univie.ac.at/ [13]). The coverage tracks revealed that the pull-down samples contained both intronic and exonic reads indicating the presence of *Tnf* and *Cxcl2* pre-mRNA, in addition to mRNA, in the TTP-bound RNA pool.

**Figure 4.**
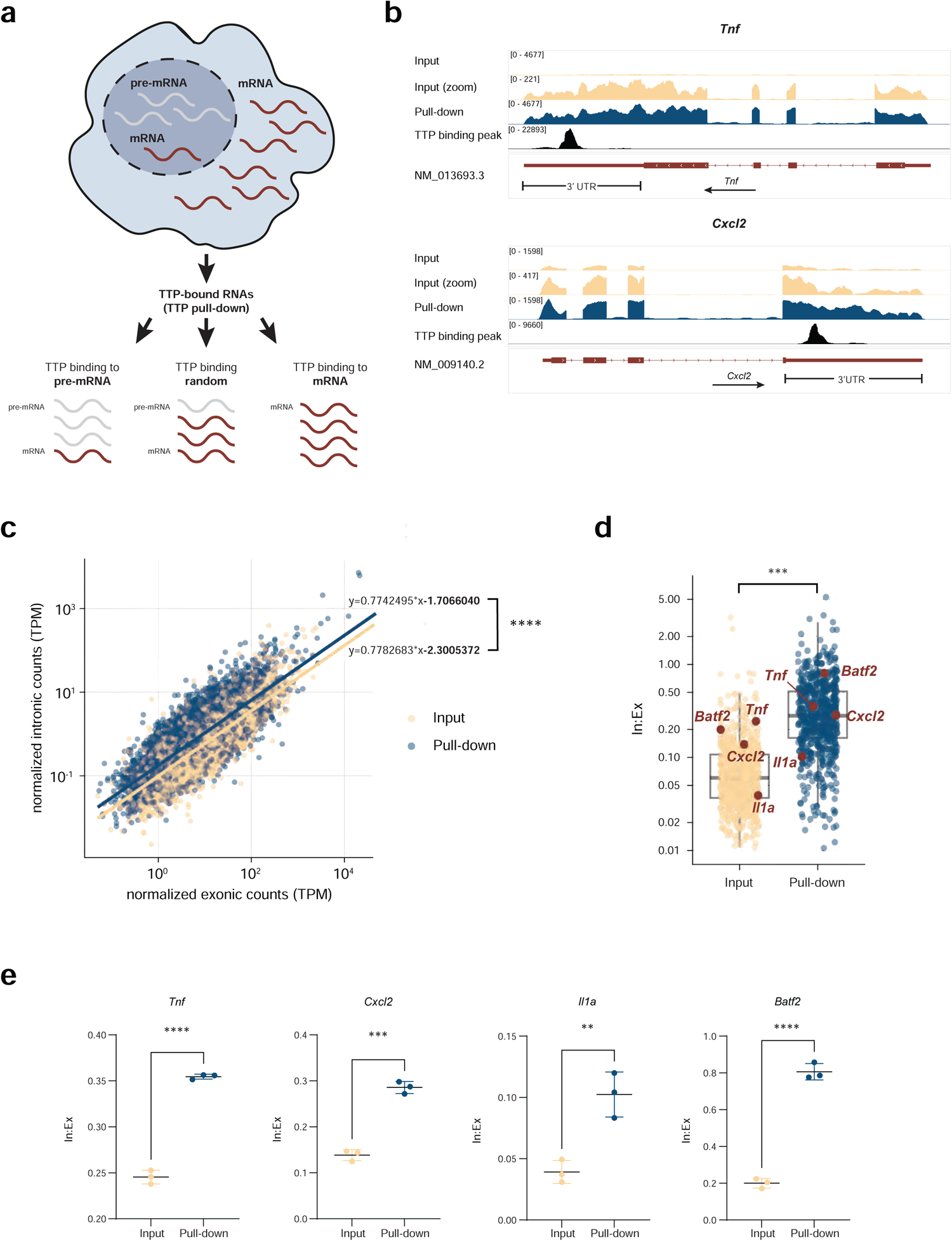
Quantitation of pre-mRNA and mRNA in TTP-bound RNA molecules in RAW macrophages. **a.** Design of pull-down experiment to test preferential binding of TTP to pre-mRNA or mRNA, indiscriminate binding to pre-mRNA and mRNA. Following pull-down of TTP-bound RNA and subsequent RNA sequencing, the pre-mRNA:mRNA ratio in pull-down sample is compared to that in input sample. **b.** Coverage plots of RNA-seq analysis of TTP-bound RNA (pull-down samples) and of total input RNA (input samples) for TTP targets *Tnf* and *Cxcl2*. TTP expression (C-terminal Myc-tagged TTP) in RAW macrophages was induced with doxycycline (0.5 µg/ml, 24 h). Subsequently, cells were treated with LPS (10 ng/ml, 6 h) and 4sU (100 µM) for 1 h (starting 5 h after LPS addition). TTP-bound RNA (Myc pull-down) and total cellular RNA (input; 10% of cell lysate used) were subjected to RNA-seq and the read coverage for *Tnf* and *Cxcl2* was visualized using Integrative Genomics Viewer (IGV). Shown are representative IGV tracks in input samples, in input with re-scaled y axis (zoom), and in pull-down samples. The rescaled y axis (zoom) allows better visualization of intronic reads (i.e. less abundant reads). The position of previously reported TTP binding peaks [13] is depicted. **c.** Visualization of mRNA counts (normalized exonic counts, x axis) and pre-mRNA counts (normalized intronic counts, y-axis) in input and pull-down samples for all detected genes. Following RNA-seq, read counts of three replicates were normalized, means for each gene were calculated and the values were plotted (input = yellow, pull-down = blue). Linear regression curves revealed significant y-intercept difference between input and pull-down samples and no difference in slopes. COvariance (ANCOVA), R function “Anova”-package “car”, n = 3, ****padj. < 0.0001. **d.** Ratios of intronic:exonic (In:Ex) counts for TTP target genes in input and pull-down samples. Example TTP target genes are depicted. Difference between In:Ex ratio in input vs. pull-down was calculated. Unpaired Student’s t test. ***padj. < 0.001. **e.** In:Ex ratio in input and pull-down samples of TTP target genes *Tnf*, *Cxcl2*, *Il1a*, and *Batf2* (means with SD, unpaired Student’s t test. **padj. < 0.01, ***padj. < 0.001, ****padj. < 0.0001).

Quantitation of pre-mRNA and mRNA was carried out by separately counting intronic and exonic reads, respectively, as described previously [35], with modifications ensuring read mapping to unambiguous intronic and exonic sequences (see Methods). PCA of normalized intronic and exonic counts in pull-down and input showed distinct clusters of corresponding replicates (**Extended Data Fig. 4a**). The means of the three replicates were then calculated for each gene detected in both, the input and pull-down RNA-seq samples (**Supplementary Table 8**), and the mean intronic counts were plotted against the mean exonic counts followed by calculation of linear regression curves (**Fig. 4c**). The regression curve intercept was significantly higher in pull-down than in input (−1.7 versus −2.3, respectively) revealing that the intronic counts were in general higher in pull-down than in input (**Fig. 4c**). The slopes of the regression curves were similar in pull-down and input (0.774 versus 0.778, respectively), showing that the intronic RNA enrichment in pull-down as compared to input was consistent across the entire spectrum of expressed genes.

Despite using the selective 4sU-facilited cross-linking approach, a contamination of the pull-down fractions with non-TTP-target RNAs could not be excluded. To determine the intronic RNA enrichment in the pull-down specifically for TTP target RNAs, we defined the TTP target gene subset (see Methods) and subsequently calculated the intron:exon (In:Ex) ratios for pull-down and input samples (**Supplementary Table 9**). A distribution analysis showed that most TTP target genes exhibited low In:Ex ratios (< 0.05) in input samples (**Extended Data Fig. 4b**), consistent with an excess of mRNA over pre-mRNA typical for total cellular RNA. Remarkably, the In:Ex ratio distribution was different in pull-down samples in that it was shifted to higher values with peaking at the In:Ex ratio of ∼0.2 (**Extended Data Fig. 4b**). This distribution analysis indicated that intronic reads were enriched in pull-down as compared to input in TTP target genes.

To quantitate this enrichment, we calculated the In:Ex ratio across all TTP target genes. Strikingly, the result revealed a significantly higher In:Ex ratio in pull-down samples as compared to input (**Fig. 4d**). The difference in the overall In:Ex ratio was well reflected at the level of individual genes as illustrated by the significantly higher In:Ex ratios for *Tnf* and *Cxcl2* in pull-down as compared to input (**Fig. 4e**). The *Tnf* and *Cxcl2* RNAs are bound by TTP in their 3’ UTRs, i.e. in exons not introns (**Fig. 4b**) (https://ttp-atlas.univie.ac.at/) [13], implying that the TTP binding sites in *Tnf* and *Cxcl2* are present in both mRNA and pre-mRNA molecules. Importantly, the TTP-bound (i.e. pull-down) *Tnf* and *Cxcl2* RNA showed a higher In:Ex ratio as compared to input RNA, thereby corroborating the binding preference of TTP for pre-mRNA rather than mRNA. If TTP did not discriminate between mRNA and pre-mRNA, the In:Ex ratios for *Tnf* and *Cxcl2* would be comparable in pull-down and input. If TTP preferred mRNA to pre-mRNA, the In:Ex ratio for *Tnf* and *Cxcl2* would be lower in pull down than in input; in this case, the In:Ex values in pull-down would likely be near zero given the much higher amounts of mRNA over pre-mRNA present in cells. Together, the enrichment of *Tnf* and *Cxcl2* pre-mRNA in pull-down samples confirmed our hypothesis that TTP preferentially binds pre-mRNA not mRNA.

A similar pre-mRNA enrichment (i.e. In:Ex higher in pull-down than in input) was observed for *Il1a*, which is a target with a dominant TTP binding peak in the 3’ UTR and a distinct TTP binding site in an intron (**Fig. 4e**) (**Extended Data Fig. 4a**). The In:Ex ratio was high (often close to 1) in targets bound by TTP predominantly in introns as exemplified by *Batf2*, which was pulled down largely as an entire pre-mRNA (**Fig. 4d**, **Extended Data Fig. 4c**). The pre-mRNA enrichment in pull-down seen in RNA-seq data was confirmed by RT-qPCR (**Extended Data Fig. 4d**). Collectively, the data demonstrate the primary target of TTP is pre-mRNA not mRNA.

### Control of inflammation through cytoplasmic mRNA decay is licensed by TTP in the cell nucleus

The binding preference of TTP for pre-mRNA not mRNA suggests that the principal place of TTP binding to a target RNA molecule is the cell nucleus. This would imply that TTP requires nuclear entry for its interaction with the target RNA. To test this hypothesis, we generated a nuclear import-deficient TTP (nuclear localization signal-mutated TTP, TTP-NLS^mut^) by introducing the mutations R126A and R130A (murine TTP coordinates) (**Extended Data Fig. 5a**). These arginine residues were reported to be essential for nuclear import of a TTP TZF peptide [36]. The R126A/ R130A double mutation prevents nuclear import also in the context of the full length TTP since HEK293 cells expressing TTP-NLS^mut^ were devoid of nuclear TTP (**Fig. 5a - c**). Cells expressing TTP-WT contained a distinct nuclear and a major cytoplasmic TTP pool, as expected (**Fig. 5a - c**). Remarkably, TTP-NLS^mut^ did not co-localize with DCP1A and EDC4 decapping foci (**Fig. 5d, e**).

**Figure 5.**
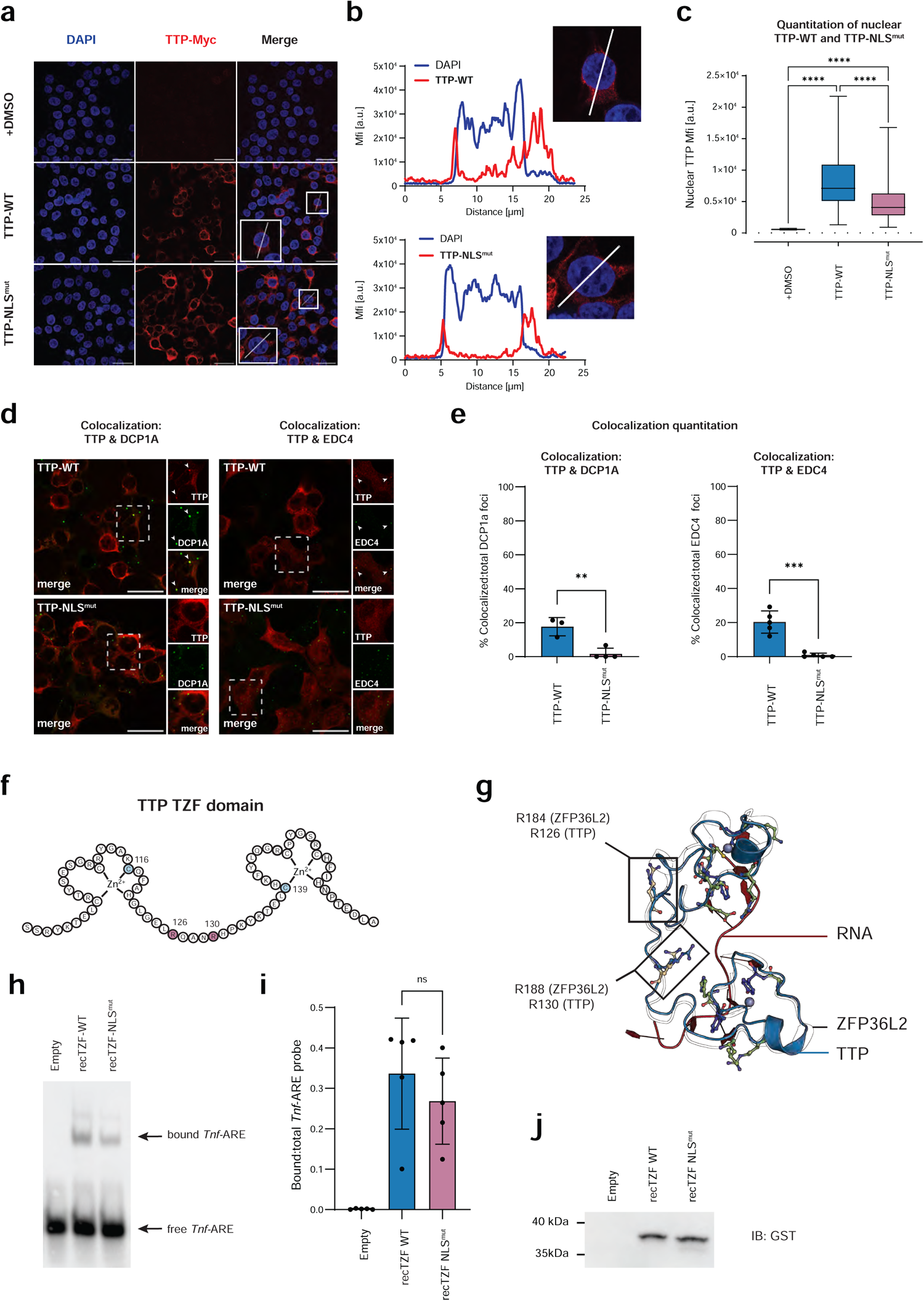
Structural modeling and RNA binding of nuclear import-deficient TTP (TTP- NLS^mut^) **a – c.** Analysis of subcellular localization of C-terminally Myc-tagged nuclear import-deficient TTP (nuclear localization signal-mutated TTP, TTP-NLS^mut^) in HEK293 cells by immunofluorescence imaging. TTP-NLS^mut^ and TTP-WT expression was induced by doxycycline (1 µg/µl, 3.5 h); control cells were treated with vehicle (DMSO). TTP constructs were detected with a Myc antibody; DAPI visualized the nucleus. (**a**) Immunofluorescence images with profile lines drawn through representative cells (enlarged 2-fold in insets) for quantitation of mean fluorescence intensity (MFI) in (b). Scale bar = 25 µm. (**b**) Quantitation of MFI along a profile line drawn through representative cells in (a). (**c**) Quantitation of nuclear MFI of TTP-NLS^mut^ and TTP-WT. Nuclear masks for quantitation of nuclear MFI were computationally defined by using ImageJ. Box represents interquartile range, horizontal line in box depicts mean. One-way analysis of variance (ANOVA) with Tukey’s multiple comparisons test, n = 62, ****padj. < 0.0001. **d, e.** Colocalization of TTP-WT or TTP-NLS^mut^ with DCP1A and EDC4 foci. TTP-WT and TTP-NLS^mut^ were expressed as in (a). (**d**) Immunofluorescence images showing TTP-WT or TTP-NLS^mut^, and endogenous DCP1A and EDC4. Large images depict TTP constructs (Myc antibody, red); highlighted cells are shown in insets together with DCP1A staining or EDC4 staining (both green) in individual and merged channels. Arrowheads point to foci of DCP1A or EDC4 colocalizing with TTP signal. Scale bar = 25 µm. (**e**) Quantitation of colocalization of TTP-WT or TTP-NLS^mut^ signals with DCP1A or EDC4 foci (Fiji plugin Comdet). Student’s t test. **padj. < 0.01, n = 44 −132 cells. **f.** Schematic representation of murine TTP TZF domain used for recombinant protein expression in *E. coli*. TZF domain corresponds to a synthetic peptide previously used in binding studies [58]; linker region amino acids R126 and R130 mutated in TTP-NLS^mut^ highlighted in lilac; zinc finger (ZF) 1 amino acid C116 and ZF2 C139 mutated in TTP-TZF^mut^ highlighted in blue. **g.** Superposition cartoon of NMR structure of ZFP36L2 TZF (black outlines) in complex with RNA (red) (PDB entry 1RGO) and AlphaFold2 model of TTP TZF (blue). ZFP36L2 TZF NMR entry is lacking the most N-terminal and 3 C-terminal amino acids in comparison to TTP TZF selected for recombinant protein studies. R184 (R126 in TTP) and R188 (R130 in TTP) shown as balls & sticks (yellow) and highlighted in rectangles. Zn ions shown as grey spheres; residues in ZFP36L2 involved in π-π staking with RNA shown in blue and residues involved in hydrogen bonds shown in green. **h, i.** Binding of recombinant TZF-WT and TZF-NLS^mut^ GST fusion proteins (recTZF-WT and recTZF-NLS^mut^) to *Tnf*-ARE assessed by RNA EMSA. (**h**) recTZF-WT and recTZF-NLS^mut^ were expressed in *E. coli* and cleared lysates were used for chemiluminescence-based RNA EMSA. *E. coli* lysates without recombinant TZF constructs were used as control (Empty). Representative RNA EMSA of 5 independent experiments is shown. (**i**) Quantitation of RNA EMSA shown in (h). Images were acquired and quantitated using ChemiDoc (BioRad). *Tnf*-ARE-bound fraction was determined as ratio of bound versus total *Tnf*-ARE. ANOVA, means with SDs, non-significant (ns), n = 5. **j.** Western blot showing expression of recTZF-WT and recTZF-NLS^mut^ in *E. coli* lysates used in (h). recTZF-WT and recTZF-NLS^mut^ was detected by immunoblotting with GST antibody (IB: GST).

The two arginines mutated in TTP-NLS^mut^ are located in the linker region connecting the two zinc fingers within the TZF domain, i.e. in the central part of TTP (**Fig. 5f**). Given that the interface for the interaction with the decapping complex is located in the N-terminal domain of TTP [27, 28], it is unlikely that the arginine mutations directly impair the interaction of TTP-NLS^mut^ with DCP1A and EDC4. Moreover, the N-terminal domain was shown to autonomously bind to the decapping proteins [27, 28]. Considering the requirement for RNA binding in association of TTP with the decapping complexes (**Fig. 2 and 3**), we reasoned that the inability of TTP-NLS^mut^ to associate with DCP1A and EDC4 foci might be caused by an impaired interaction with target RNA in cells. We first assessed the potential of the R126A/R130A double mutation in TTP-NLS^mut^ to directly affect RNA binding by structural modeling of the TZF domain. As no structure of full length TTP or the TTP TZF is available, we employed the reported NMR structure of the TZF of the TTP homolog ZFP36L2 in complex with RNA (PDB entry 1RGO) [37]. The TZF domain is conserved in the TTP protein family (**Extended Data Fig. 3a**) and is expected to acquire a similar structure in all TTP protein family members. This expectation was supported by the comparison of the reported NMR structure of the first zinc finger of TTP (PDB entry 1M9O) [38] with the ZFP36L2 TZF (PDB entry 1RGO) (**Extended Data Fig. 5b**).

On the grounds of these similarities, we assessed the structural role of the arginine residues that are mutated in TTP-NLS^mut^, and the potential effects of the R126A/R130A mutations on the zinc fingers and RNA binding. The analysis of conformations of the critical arginines within the respective NMR ensembles of ZFP36L2 TZF and TTP zinc finger 1 structural data revealed a high flexibility of these arginines (**Extended Data Fig. 5b**). Moreover, there is no direct interaction between either of the two relevant arginines and the bound RNA molecule in the ZFP36L2 TZF NMR ensemble (**Fig. 5g**). AlphaFold2 modeling predicted the overall fold of the TTP TZF to be highly similar to that of the ZFP36L2 TZF (**Fig. 5g**), with the critical arginine residues positioned within the conformational space observed in the NMR structures. Consequently, interactions of the two TTP TZF arginines with RNA are unlikely.

To further assess the potential effect of the arginine mutations in TTP-NLS^mut^ on the overall TZF fold we performed structure predictions with AlphaFold2 using CoLabFold, and comparative modelling using the Robetta server. The structural models indicated that the introduction of the smaller alanine residues, which cannot form hydrogen bonds, may increase the flexibility of the linker region as seen from the increased TTP-NLS^mut^ to ZFP36L2 root-mean-square-deviation (RMSD) compared to that of TTP-WT to ZFP36L2 (**Supplementary Table 10**). However, as the main interaction interfaces of TTP and ZFP36L2 with RNA are the zinc fingers it is unlikely that an increased linker flexibility has a significant impact on the RNA bound state (**Extended Data Fig. 5c**). Collectively, the structural analyses focusing on the TZF domain indicate that the arginine residues R126 and R130 do not contribute to RNA binding, and that the mutation of these residues to alanine does not affect the TZF structural integrity.

To address the possible effect of the R126A/R130A double mutation on RNA binding experimentally, we expressed the WT and mutated TZF domains as GST fusion proteins in *E. coli* and used these recombinant proteins (recTZF-WT and recTZF-NLS^mut^) in RNA EMSA experiments. The results showed that both recTZF-WT and recTZF-NLS^mut^ were able to bind to RNA although the binding of recTZF-NLS^mut^ was weaker when compared to recTZF-WT (**Fig. 5i**). A titration experiment using unlabeled *Tnf*-ARE in various ratios to the labeled *Tnf* ARE verified that the labeled *Tnf*-ARE was in a similar excess in both, the recTZF-WT and recTZF-NLS^mut^ EMSA experiments (**Extended Data Fig. 5d, e**).

Together, the structural TZF analysis and EMSA experiments using recombinant TZF constructs indicate that the RNA binding site remains functional in TTP-NLS^mut^. We then proceeded to test whether TTP-NLS^mut^, i.e. a TTP variant that is unable to enter the nucleus hence to interact with pre-mRNA, binds target RNAs in cells. By employing immunostimulated RAW macrophages we made sure that natural TTP target RNAs are expressed. Binding of TTP to RNA in cells was analyzed using the CLIP approach described in **Fig. 4**. We inducibly expressed TTP-WT, TTP-NLS^mut^, and TTP-TZF^mut^ in RAW macrophages (**Extended Data Fig. 6a**), treated them with LPS and 4sU, and subsequently quantitated the amounts of *Il1a*, *Tnf* and *Il6* mRNAs bound to these TTP constructs in cells. The experiment revealed that the binding of TTP-NLS^mut^ to the target mRNAs in cells was dramatically decreased, resembling the RNA binding-deficient TTP-TZF^mut^ construct (**Fig. 6a**). Consistent with the impairment in mRNA targeting by TTP-NLS^mut^ in cells, the mRNA levels of *Il1a*, *Tnf* and *Il6*, which all code for key pro-inflammatory cytokines, were higher in RAW macrophages expressing TTP-NLS^mut^ as compared to TTP-WT cells (**Fig. 6b**). The deficiency of TTP-NLS^mut^ to control *Il1a*, *Tnf* and *Il6* mRNA levels mimicked the properties of TTP-TZF^mut^ (**Fig. 6b**). These experiments suggest that the impaired association of TTP-NLS^mut^ with the DCP1a and EDC4 foci results from a failure of the nuclear import-deficient TTP mutant to interact with RNA in cells despite having a functional RNA-binding domain.

**Figure 6.**
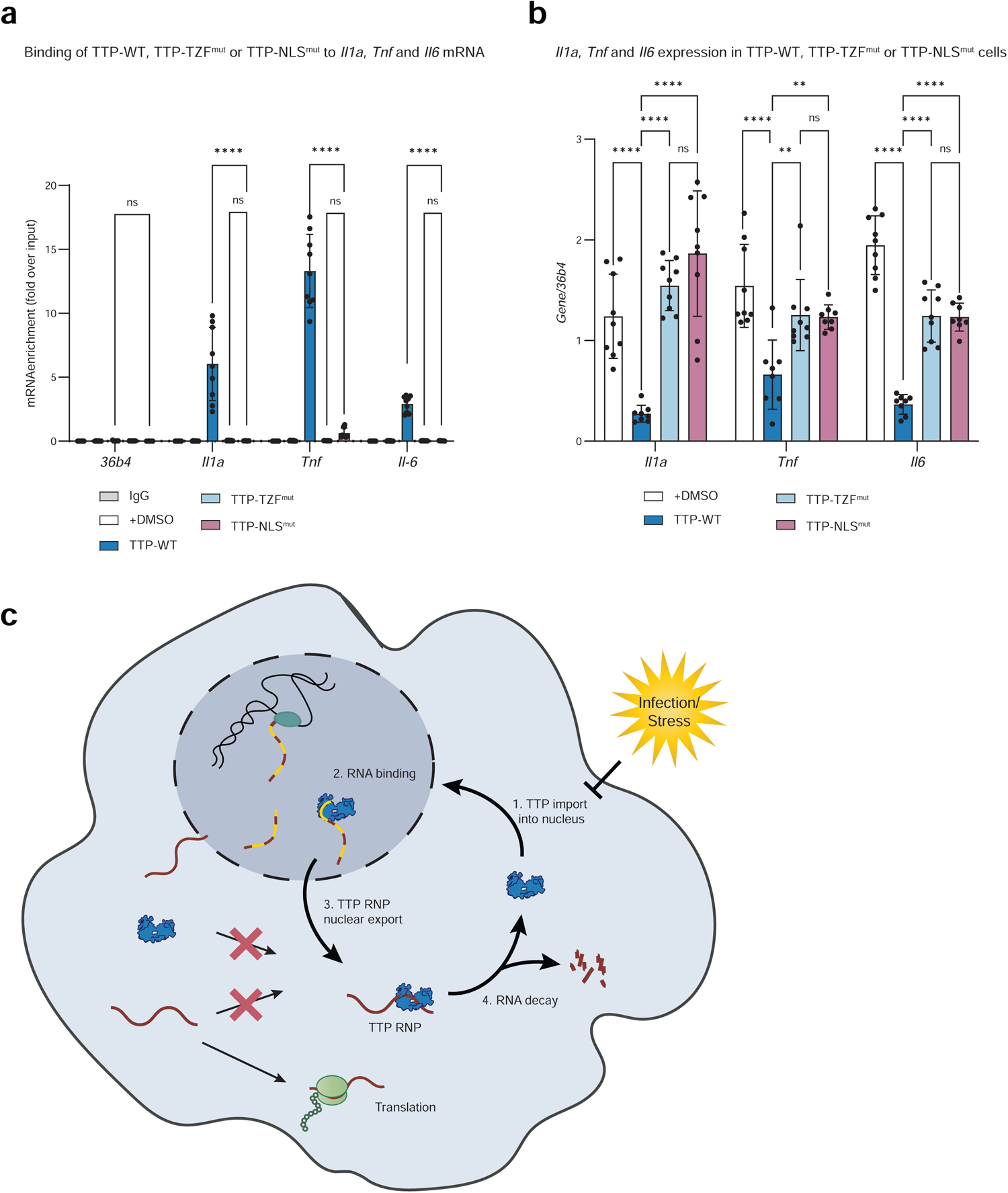
Analysis of de novo binding of TTP to mature cytoplasmic mRNA in immunostimulated RAW macrophages. **a.** qRT-PCR analysis of mRNAs bound to TTP (TTP-WT) or to constitutively cytoplasmic TTP (TTP-NLS^mut^). TTP-WT and TTP-NLS^mut^ were inducibly expressed in RAW macrophages as described in Figure 3h. TTP-TZF^mut^ was included as negative control. Cells were treated with LPS and 4sU as described in Extended Data Figure 4d. TTP-bound RNA (Myc pull-down) and total cellular RNA (input; 10% of cell lysate used) were assessed by qRT-PCR. Controls: mouse IgG isotype antibody instead of Myc antibody, and RAW macrophages treated with vehicle (DMSO) instead of doxycycline. Enrichment of *Il1a, Tnf* and *Il6* mRNA in pull-down fraction from TTP-WT, TTP-NLS^mut^, and TTP-TZF^mut^ RAW macrophages is shown as fold difference between pull-down and input. *36b4* mRNA was used and non-TTP target control. One-way analysis of variance (ANOVA), means of RT-qPCR data with SDs, n = 3 (each in triplicates), ***padj. < 0.001, ****padj. < 0.0001. **b.** Levels of *Il1a, Tnf* and *Il6* mRNA in RAW macrophages inducibly expressing TTP-WT, TTP-NLS^mut^, and TTP-TZF^mut^ constructs assessed by qRT-PCR. Expression of TTP constructs was induced with doxycycline (0.5 µg/ml, 24 h) followed by LPS stimulation (10 ng/ml, 6 h). Note that RAW macrophages were devoid of endogenous TTP as in all other experiments in this study. Expression of the housekeeping gene *36b4* was used for normalization. Data are presented as means and SDs, with all data points shown. ANOVA n = 3, each measured in triplicates, ***padj. < 0.001, ****padj. < 0.0001. **c.** Model of mRNA degradation by a TTP binding & nucleo-cytoplasmic shuttling cycle. The model proposes that, following TTP translation or release upon degradation of bound mRNA, TTP translocates to the nucleus (Step 1), to bind pre-mRNA (Step 2). Subsequently, pre-mRNA is processed to mRNA, and the TTP-mRNA complex is exported to the cytoplasm (Step 3) where it associates with RNA degradation machinery to eliminate the bound RNA (Step 4). External stress cues enforce cytoplasmic localization of TTP.

Collectively, the results imply that cytoplasmic TTP molecules are unable to bind target mRNA *de novo*. Together with the documented degradation of TTP-bound RNAs in the cytoplasm [39], these data imply that TTP licenses the target degradation by binding to it in the nucleus, i.e. prior to mRNA export to the cytoplasm. This nuclear licensing mechanism is supported by our data demonstrating preferential binding of TTP to pre-mRNA not mRNA (**Fig. 4**).

## DISCUSSION

Our studies reveal that TTP prioritizes pre-mRNA over mRNA for binding while cytoplasmic mRNA does not constitute a de novo TTP target. Furthermore, we demonstrate that RNA binding prevents nuclear accumulation of TTP and enables TTP to associate with the cytoplasmic RNA degradation complexes. Importantly, the interaction of TTP with the cytoplasmic RNA degradation complexes is dependent on the ability of TTP to enter the nucleus and engage with the target RNA in this compartment. Overall, these results are consistent with a model in which TTP binds pre-mRNA in the nucleus and, following maturation of pre-mRNA to mRNA, the TTP-mRNA complex is subsequently exported to the cytoplasm (**Fig. 6c**). The cytoplasmic TTP-mRNA complex associates with the RNA degradation machinery to eliminate the bound RNA and curb inflammation (**Fig. 6c**). Furthermore, our data suggest that the RNA binding-dependent hierarchical model is conserved within the TTP protein family.

Early studies suggested that TTP associates with the decapping and deadenylation enzymes irrespectively of RNA binding and, consequently, that excessive TTP might sequester the RNA degradation complexes from other processes and thereby inhibit mRNA degradation [9–11, 22]. This model was later found to be inconsistent with the analysis of mice bearing a zinc finger mutation in the TTP locus (C116R knock-in mice), which unequivocally showed that the RNA binding-deficient TTP mutant did not exhibit any dominant-negative effects [40]. Our study provides mechanistic explanations for the results obtained with the C116R knock-in mice in that it excludes the possibility that TTP might seize the decapping and deadenylation enzymes in the RNA-unbound state. Our analysis of the proximal interactomes of TTP and ZFP36L1 indicates that the requirement for RNA binding in the interaction of these proteins with the RNA degradation complexes is conserved in the TTP protein family.

This hierarchical assembly of TTP-containing decapping and deadenylation structures is consistent with the current model of the formation of RNP granules, including P-bodies [41]. The model proposes that RNA-processing, membrane-less granules are hierarchically assembled biomolecular condensates, that are dependent upon multi-valent interactions between RNAs and proteins. Proteins with intrinsically disordered regions (IDRs) play an important role in the orchestration of the RNP granule formation [42]. In agreement, TTP was recently found to contain an N-terminal IDR, which facilitates the interaction with DCP2 [27]. Another low complexity region is located in the C-terminal part of TTP suggesting that this region might facilitate other IDR-mediated interactions such as those with the CCR4-CNOT complex, which was shown to bind the TTP C-terminal domain [10, 11, 28, 43].

TTP is a largely cytoplasmic protein and exhibits nucleo-cytoplasmic shuttling. The significance of this shuttling, however, remained unknown [24, 29, 44]. Our study reveals that TTP prioritizes pre-mRNA over mRNA for binding implicating that the translocation into the nucleus is essential for the interaction of TTP with its natural target RNA. This conclusion is supported by the impaired interaction of a nuclear import-deficient TTP mutant with endogenous target RNAs and, consequently, with the RNA degradation complexes. Together, the data propose that periodic nuclear import and export are indispensable for the TTP life cycle as they continuously couple target binding with target degradation. The predominantly cytoplasmic localization of TTP indicates that the nuclear dwell time of TTP is short. Surprisingly, the appropriate partitioning of TTP into a larger cytoplasmic and smaller nuclear pool is dependent on RNA binding since TTP-TZF^mut^ accumulates in the nucleus. We see the same role of RNA binding also in ZFP36L1 suggesting that the regulation of subcellular localization by RNA binding is conserved in the TTP protein family.

Previous studies investigating the impact of external cues on the subcellular distribution of TTP showed that stress signaling promotes cytoplasmic accumulation of TTP [24, 45]. Our proximal labeling data demonstrate that the divergent steady-state interactomes of TTP and TTP-TZF^mut^ acquire a convergent composition upon stress. Moreover, the nuclear signature found in the steady-state TTP-TZF^mut^ interactome is lost following stress induction, implying that stress signals determine the localization of TTP regardless of its binding to RNA. These results highlight the significance of stress-induced cytoplasmic accumulation of TTP since the interaction of TTP with its principal target, i.e. pre-mRNA, is efficiently prevented in this way (**Fig. 6c**).

The deficiency of TTP-NLS^mut^ to bind TTP targets in cells is intriguing given that structural modeling and assessment of RNA binding by recombinant TZF proteins did not indicate any major perturbances in recTZF-NLS^mut^ as compared to wild-type TZF, although we noticed a minor decrease in the RNA binding of the recTZF-NLS^mut^. The limitation of our structural modeling is the absence of structural data other than the TZF domain. Moreover, TTP is predicted to be unstructured and intrinsically disordered, except for the TZF domain [27]. Hence, the impact of the N-terminal and C-terminal domains on the centrally located TZF domain in cells cannot be currently estimated. Given that N-terminal and C-terminal domains constitute the protein interaction hubs of TTP and are subject to multiple phosphorylations [10, 11, 25, 28, 43, 46], it is feasible that they indirectly regulate the interaction of TTP with RNA in cells.

The interaction of TTP with pre-mRNA occurs probably early during the transcription process as evident from the high RNA-seq coverage of full-length pre-mRNAs bound to TTP, irrespective of whether the binding sites are in 3’ UTRs or introns. The biological significance of TTP binding to intronic regions remains unclear, nevertheless our data suggest that such binding might increase TTP crowding in the nucleus and thereby promote the nuclear dwell time of TTP.

The importance of precise regulation of immune responses is underlined by the multilayered control of essentially every step of the immune response at the cellular and organismal level. Our study unravels a nucleus-based fate decision mechanism ensuring that mRNAs encoding pro-inflammatory mediators are targeted for degradation prior to the pioneering round of translation. This mechanism is driven by binding of TTP to pre-mRNA which, following pre-mRNA processing, licenses the target for subsequent degradation in the cytoplasm. The proposed model minimizes the risks of spurious inflammation-promoting translation that would occur if stochastic processes drove mRNA decay during the immune response.

## METHODS

### Plasmids

TTP, TTP-TZF C116R/C139R double point mutant (TTP-TZF^mut^) and ZFP36L1 (from murine colon total RNA) were cloned into a modified, doxycycline hyclate (dox, Sigma-Aldrich, D9891)-inducible lentiviral expression vector pCW 57.1 (Addgene #41393) by Gibson assembly. In brief, C-terminal 3xMyc-tag and backbone complementary overhangs for Gibson assembly were added by PCR, purified using column DNA Clean-up kit (Monarch PCR & DNA Cleanup Kit, # T1030L, NEB) and mixed with Nhe1 and BamH1 double-digested and gel-purified target pCW57.1 vector in a molecular ratio of 1:2. Gibson assembly was performed following the manufacturer’s instruction (NEB, E2621S) and transformed into chemically competent DH10B cells. Lentiviral C-terminally 3xMyc-tagged-TTP NLS point mutant (TTP-NLS^mut^) was generated by mutating R126 and R130 to alanine using PCR-based site-directed mutagenesis of the parental pCW57.1_tetON_TTP-3xMyc. GFP was PCR-amplified from pEGFP-N1 (Clontech) and used for Gibson cloning. APEX2 constructs were obtained by cloning of TTP-WT, TTP-TZF^mut^, ZFP36L1-WT, ZFP36L1-TZF^mut^ and GFP into a modified pCW57.1 lentiviral expression vector containing APEX2. All expression constructs contained a puromycin resistance cassette for selection. N-terminally TurboID TTP (TID-TTP) and empty TurboID (TID-E) vectors were generated by cloning TID from V5-TurboID-NES_pCDNA3 (Addgene#107169) into a modified TurboID-containing pCW57.1 vector (pCW57.1-pCW 57.1-MYC-TurboID-MCS-PGK-mCherry-P2A-rtTA), expressing a constitutively-active mCherry selection cassette. TID-TTP-TZF^mut^ was generated by exchanging TTP-WT in parental vector with TTP-TZF^mut^ by conventional restriction digest cloning. All constructs were verified by sequencing.

### Cell culture and lentiviral transduction

All cell lines were cultured in DMEM (Sigma-Aldrich, D6429) containing 10% fetal bovine serum (FBS) and 1% penicillin/streptomycin at 37°C and 5% CO2 in a humified incubator. RAW 264.7 macrophages used in all experiments bore a CRISPR/Cas9-generated deletion in the TTP gene. Cells were seeded 24 h prior to treatments which were directly added to cell culture dishes without change of the cell culture medium. Lentiviral particles were generated by co-transfecting 1.5 µg of total DNA of pMDLg/pRRE (Addgene #12251), pCMV-VSV-G (Addgene #8454), and pRSV-Rev (Addgene #12253) with modified pCW57.1 target vector in a ratio of 5 : 1 : 5 : 5 into HEK293 cells using polyethyleneimine (PEI, Polysciences, 23966). Three days post-transfection, supernatants containing viral particles were harvested, passed through a 0.45 µm filter and used to infect either HEK293 cells in a dilution of 1:5 or RAW macrophages in a dilution of 1:2 with 8 µg/ml polybrene (Sigma-Aldrich, TR1003G). Transduced cells were selected by either puromycin (Sigma-Aldrich, P8833-100MG) (3 µg/ml for HEK293 cells or 8 µg/ml for RAW macrophages) for 1 week, or by FACS for mCherry-positive cells on day 3 post-infection.

### Cell lysates and Western blotting

Expression of TTP-constructs in HEK293 cells and RAW macrophages was induced by 1 µg/µl for 3.5 h (HEK293 cells) or 0.5 µg/ml for 24 h (RAW macrophages). After induction, cells were washed twice with PBS and lysed in either Frackelton buffer (10 mM Tris-HCl pH8, 30 mM Na4P2O7, 50 mM NaCl, 50 mM NaF, 1% Triton X-100 and 1x cOmplete protease inhibitor cocktail) or RIPA buffer (50mM Tris HCl pH8, 150mM NaCl, 0.1% SDS, 0.5% sodium deoxycholate, 1% NP-40, 1mM PMSF, 1x cOmplete protease inhibitor cocktail) for 5 min on ice. Lysates were cleared by centrifugation at 13200 rpm for 5 min at 4°C and supernatant was transferred into a fresh tube. Protein concentration was determined using Pierce BCA Protein Assay Kit (Thermo-Fisher Scientific, Cat #23225). After concentration adjustment, lysates were mixed at a ratio of 2:1 with SDS loading dye and heated for 5 min at 95°C. Samples were loaded onto 10% SDS polyacrylamide gel for separation, followed by wet transfer onto nitrocellulose membrane (GE Healthcare) in carbonate transfer buffer (3 mM Na_2_CO_3_, 10 mM NaHCO_3_, and 20% ethanol) for 16 h at 200 mA and additional 2 h at 400 mA at 4°C. Membranes were blocked in 10% BSA in TBS-T (20 mM Tris, 150 mM NaCl, 0.5% Tween 20, pH 8.0) for 1 h at room temperature (RT) and subsequently incubated with primary antibody overnight at 4°C. Membranes were then washed three times with TBS-T for 5 min each, and incubated with HRP-conjugated secondary antibody for 1 h at RT. Chemiluminescence was detected and quantified by using a ChemiDoc Touch Imaging System (BioRad).

### Immunofluorescence

2×10^4^ HEK293 cells, expressing C-terminally 3x Myc-tagged TTP or ZFP36L1 were seeded onto poly-lysine (Sigma-Aldrich, P4832) pretreated glass coverslips for 48 h. TTP or ZFP36L1 expression was induced by doxycycline (1 µg/ml) for 3.5 h. Cells were then washed three times with PBS, fixed in 4% formaldehyde in PBS for 10 min at RT, washed three times with PBS, quenched with 0.1 M glycine in PBS for 10 min on RT, washed three times with PBS and subsequently permeabilized with 0.3% Triton in 3% BSA (w/v) in PBS for 30 min on RT. Primary antibodies (Myc 1:2000, EDC4 1:500, DCP1a 1:500) were diluted in permeabilization solution and incubated for 1 h at RT. After three PBS washes, slides were incubated with secondary antibodies conjugated to Alexa Fluor 594 (for Myc) or Alexa Fluor 488 (for EDC4 and DCP1a) (Thermo-Fisher Scientific, Cat # A-11005 and A-11008) for 1 h at RT. Samples were washed three times with PBS and nuclei were stained with DAPI (0.5 µg/ml, Sigma-Aldrich, D8417-10MG) for 10 min on RT. Samples were then washed twice with PBS and once with H_2_O and subsequently mounted onto microscopy slides with Prolong Gold antifade reagent (Thermo-Fisher Scientific, P36934). Confocal images were captured on the Zeiss LSM 980 microscope using a 64x oil objective (Plan-Apochromat 1.4NA Oil) with a PMT detector at RT with Zeiss ZEN 3.3 software. All images were analyzed in the Fiji distribution of ImageJ [47]. The mean fluorescence intensity (MFI) of nuclear TTP was measured by segmenting cells using Otsu thresholding in the DAPI channel to define nuclei boundaries as regions of interest (ROI). The MFI of ROIs was then measured in the Alexa Fluor 594 (Myc) channel. Profile plots were generated with the plot profile function in Fiji by drawing a line through a cell of interest. Data generated by the plot profile function were transferred to Graphpad to obtain profile plots depicting the intensity profiles of respective channels along the line. Colocalization analysis was performed using Comdet v.0.5.5 plug-in in Fiji to detect colocalizing particles in Alexa Fluor 594 (Myc) and Alexa Fluor 488 (EDC4, DCP1a) channels. Maximum distance between colocalized spots was set to 4 pixels (∼0.28 µm). In Alexa Fluor 594 (Myc) channel particles had to reach a size of 7 pixels and an intensity threshold of 50-75, in Alexa Fluor 488 channel particles had to reach a size of 10 pixels and an intensity threshold of 100-200.

### APEX2 proximity labeling

APEX2 (APX) proximity labeling in HEK293 cells was largely performed as described previously [48] with the following adjustments. Lentivirally-transduced bait protein expression was induced in confluent 15 cm tissue culture dish by 1 µg/ml dox for 3.5 h for TTP-APX experiments. For ZFP36L1 labeling experiments, bait protein expression was induced for 24 h using doxycycline at 0.02-1 µg/ml to obtain similar expression levels. Where indicated, samples were treated with 100 µM anisomycin for 1.5 h prior to labeling. Biotin-phenol was added to cells for 30 min (500 µM final concentration) prior to labeling. Labeling was induced by treating cells with 1 mM H_2_O_2_ for 1 min. Subsequently, cells were quenched by washing three times with quenching medium (10 mM sodium ascorbate, 5 mM Trolox and 10 mM sodium azide in PBS) and three times with PBS. Labeled cells were lysed with RIPA buffer, and protein concentration was measured using Pierce Protein Assay Kit. Prior to affinity purification, streptavidin bead slurry (Thermo-Fisher Scientific, Cat #88817) was acetylated with 5 mM sulfo-NHS acetate in 50 mM ammonium bicarbonate with 0.2% Tween at RT for 1 h, as described [49]. In brief, beads were washed three times with 50 mM HEPES, pH 7.8, 0.2% Tween on a magnetic rack, incubated with sulfo-NHS acetate while rotating for 1 h at RT, subsequently washed in 50 mM ammonium bicarbonate, 0.2% Tween, and finally stored in PBS-T (0.2% Tween), 0.02% sodium azide at 4°C until further use. Total protein lysate (1 mg) was loaded onto 150 µl acetylated streptavidin bead slurry and incubated overnight at 4°C while rotating. Samples were then washed to remove unspecific binders. Additional washes with 50 mM HEPES pH 7 (5 - 7 times) were included prior to mass spectrometry submission to remove residual detergents. For Western blot analysis, 10% input sample, flow-through samples or eluted samples (pull-down) were taken. Elution of pull-down was facilitated by heating the bead slurry in 1x SDS loading dye at 95°C for 5 min.

### TID proximity labeling

TID proximity labeling in RAW macrophages was performed similarly to APEX2-based labeling in HEK293 cells with the following differences. Lentivirally transduced RAW macrophages were sorted for similar mCherry signal by FACS and subsequently seeded into 15 cm tissue culture dishes (7×10^6^ cells per dish) 48 h prior to the experiment. TID-protein expression was induced the next day with 0.025 µg/ml dox (TID-E) or 0.5 µg/ml dox for 24 h to achieve similar expression of constructs. Labeling was performed by supplying cells with 500 µM biotin for 15 min. Afterwards, cells, were washed three times with PBS and lysed in RIPA lysis buffer (see above). 2 mg of cell lysates were incubated with 100 µl acetylated streptavidin bead slurry overnight at 4°C while rotating. Washing steps and sample preparation for mass spectrometry submission as well as Western blot analysis were identical to APEX2 labeling in HEK293 cells.

### Sample preparation for mass spectrometry analysis

Beads were transferred to new tubes and resuspended in 50 µl of 50 mM ammonium bicarbonate (ABC). Disulfide bonds were reduced with 10 mM dithiothreitol for 30 min at RT before adding 25 mM iodoacetamide and incubating for another 30 min at RT in the dark. The remaining iodoacetamide was quenched by adding 5 mM DTT and the proteins were digested with 150 ng LysC (mass spectrometry grade, FUJIFILM Wako chemicals) in 1.5 µl 50 mM ammonium bicarbonate at 25°C overnight. The supernatant without beads was incubated with 150 ng trypsin (Trypsin Gold, Promega) in 1.5 µl 50 mM ammonium bicarbonate at 37°C for 5 h. The digest was stopped by the addition of trifluoroacetic acid (TFA) to a final concentration of 0.5 %, and the peptides were desalted using C18 Stagetips [50].

### Liquid chromatography-mass spectrometry analysis

Peptides were separated on an Ultimate 3000 RSLC nano-flow chromatography system (Thermo-Fisher Scientific), using a pre-column for sample loading (Acclaim PepMap C18, 2 cm × 0.1 mm, 5 μm, Thermo-Fisher Scientific), and a C18 analytical column (Acclaim PepMap C18, 50 cm × 0.75 mm, 2 μm, Thermo-Fisher Scientific), applying a segmented linear gradient from 2% to 35% and finally 80% solvent B (80 % acetonitrile, 0.1 % formic acid; solvent A 0.1 % formic acid) at a flow rate of 230 nl/min over 120 min.

In TTP-APX labeling experiment, eluted peptides were analyzed on a Q Exactive HF-X Orbitrap mass spectrometer (Thermo-Fisher Scientific), which was coupled to the column with a customized nano-spray EASY-Spray ion source (Thermo-Fisher Scientific) using coated emitter tips (New Objective). The mass spectrometer was operated in data-dependent acquisition mode (DDA), and survey scans were obtained in a mass range of 375-1500 m/z with lock mass activated, at a resolution of 120k at 200 m/z and an AGC target value of 3E6. The 25 most intense ions were selected with an isolation width of 1.4 m/z, isolation offset 0.0 m/z, fragmented in the HCD cell at 28% collision energy and the spectra recorded for max. 100 ms at a target value of 1E5 and a resolution of 30k. Peptides with a charge of +1 or >+6 were excluded from fragmentation, the peptide match feature was set to preferred, the exclude isotope feature was enabled, and selected precursors were dynamically excluded from repeated sampling for 30 seconds.

In L1-APX and TID-TTP labeling experiments, eluting peptides were analyzed on an Exploris 480 Orbitrap mass spectrometer (Thermo-Fisher Scientific) coupled to the column with a FAIMS pro ion-source (Thermo-Fisher Scientific) using coated emitter tips (PepSep, MSWil) with the following settings: The mass spectrometer was operated in DDA mode with two FAIMS compensation voltages (CV) set to −45 or −60 and 1.5 s cycle time per CV. The survey scans were obtained in a mass range of 350-1500 m/z, at a resolution of 60k at 200 m/z, and a normalized AGC target at 100%. The most intense ions were selected with an isolation width of 1.2 m/z, fragmented in the HCD cell at 28% collision energy, and the spectra recorded for max. 100 ms at a normalized AGC target of 100% and a resolution of 15k. Peptides with a charge of +2 to +6 were included for fragmentation, the peptide match feature was set to preferred, the exclude isotope feature was enabled, and selected precursors were dynamically excluded from repeated sampling for 45 seconds.

### Proteomics data analysis

MS raw data split for each CV (FreeStyle 1.7, Thermo-Fisher Scientific) were analyzed using the MaxQuant [51] (version 1.6.17.0 for TTP-APX experiment and TID-TTP experiment, version 2.3.1.0 for L1-APX experiment) with the Uniprot human reference proteome (version 2020.01 for TTP-APX experiment, version 2022.05 for L1-APX experiment) and Uniprot mouse reference proteome (version 2020.01 for the TID-TTP experiment), as well as a database of most common contaminants and target protein sequences (mouse TTP in TTP-APX experiments and mouse ZFP36L1 in L1-APX experiment). The search was performed with full trypsin specificity and a maximum of two missed cleavages at a protein and peptide spectrum match false discovery rate of 1%. Carbamidomethylation of cysteine residues was set as fixed, oxidation of methionine, and N-terminal acetylation as variable modifications. For label-free quantitation the “match between runs” only within the sample batch and the LFQ function were activated - all other parameters were left at default.

MaxQuant output tables were further processed in R 4.2.1 (https://www.R-project.org) using amica (TTP-APX dataset) or Cassiopeia_LFQ (all other datasets) (https://github.com/maxperutzlabs-ms/Cassiopeia_LFQ). Reverse database identifications, contaminant proteins, protein groups identified only by a modified peptide, protein groups with less than two quantitative values in one experimental group, and protein groups with less than 2 razor peptides were removed for further analysis. Missing values were replaced by randomly drawing data points from a normal distribution model on the whole dataset (data mean shifted by −1.8 standard deviations, a width of the distribution of 0.3 standard deviations). Differences between groups were statistically evaluated using the LIMMA 3.52.1 [52] at 5% FDR (Benjamini-Hochberg).

The output from Cassiopeia was further analyzed with amica software [26]. Differential expression analysis was performed with DeqMS (version 1.10.0) [53]. In detail, amica was used for principal component analysis, differential analysis of peptide intensities, and over-representation analysis (ORA) using gprofiler2 (version 0.2.1) [54] (log_2_ FC > 1 and padj. ≤ 0.05). For ORA GO: CC, GO: MF, GO: BP of the gene ontology database [55] were used.

### Proteomics data deposition

The mass spectrometry proteomics data have been deposited to the ProteomeXchange Consortium via the PRIDE partner repository [56] with the dataset identifier PXD046381.

### Pull-down of TTP-bound RNA for RNA-Seq and RT-qPCR analyses

Metabolic labeling, crosslinking, harvest and immunoprecipitation of RAW macrophages for qPCR analysis or total RNA seq was performed as described previously [13], with modifications. Briefly, RAW macrophages (15×10^6^ cells) were seeded into 15 cm tissue culture treated dishes 48 h prior to crosslinking and harvesting. Expression of TTP constructs was induced with 0.5 µg/µl doxycycline for 24 h in addition to a control omitting doxycycline treatment. Cells were stimulated with 10 ng/ml LPS for 6 h. Cells were supplied with 100 µM 4sU 7 h (in qPCR experiment) or 1 h (in RNA-Seq experiment) before harvest. TTP-bound RNA was precipitated using 50 µl Pierce™ anti-c-Myc magnetic bead slurry (Thermo-Fisher Scientific, Cat #88843) for 16 h at 4 °C. For isotype control, IgG (CST, Cat #5415) bound to Protein A/G magnetic beads (Thermo-Fisher Scientific, Cat # 88802), was included. 10% of total cell lysate per sample was used as total input RNA. After immunoprecipitation, beads were washed in high salt buffer (50 mM Tris-HCl pH 7.4, 1 M NaCl, 1 mM EDTA, 1% NP-40, 0.1% SDS, 0.5% sodium deoxycholate) prior to wash buffer as described [13]. TTP-bound RNA, as well as input RNA, was purified after proteinase K digestion by acidic phenol-chloroform extraction (Ambion) and ethanol precipitation. cDNA was produced using random hexamer primers (Mycrosynth) with recombinant M-MuLV reverse transcriptase (RevertAid, Thermo-Fisher Scientific, Cat #K1691) and used for quantitation of TTP-bound RNA and total RNA by RT-qPCR with HOT FIREPol EvaGreen qPCR Supermix (Medibena, Cat #SB_08-36-GP) on CFX Touch qPCR cyclers (Bio-Rad). Relative expression levels were calculated with the CFX Maestro Analysis software, as described previously [57]. The following primers were used for qPCR: *36B4* fw TCCTTCTTCCAGGCTTTGGG and rv GGACACCCTCCAGAAAGCGA; *Il1a* fw CAAACTGATGAAGCTCGTCA and rv TCTCCTTGAGCGCTCACGAA; pre-*Il1a* fw GGGACCACTCTCACTAAGCC and rv CTTCCCGTTGCTTGACGTTG; *Il6* fw AGTTGCCTTCTTGGGACTGA and rv TTCTGCAAGTGCATCATCGT; *Tnf* fw GATCGGTCCCCAAAGGGATG and rv CACTTGGTGGTTTGCTACGAC; pre-*Tnf* fw GGCAAAGAGGAACTGTAAG and rv CCATAGAACTGATGAGAGG.

### RNA-seq

For total RNA-seq, RNA quality was assessed using Agilent RNA 6000 Nano Assays (5067-1511) on an Agilent 2100 Bioanalyzer. RNA sequencing was performed in the VBCF (Vienna BioCenter Core Facilities) Next Generation Sequencing (NGS) Facility (www.viennabiocenter.org/facilities). Library preparation, quality control and sequencing were performed by the NGS facility. Total RNA input samples were subjected to automated ribonucleic depletion using rRNA depletion kit Ribovanish prior to library preparation. Library was prepared using Ultra II Directional RNA Library Prep Kit (NEB, Cat #E7765) for all samples. After library-quality check, samples were sequenced paired end (50 bp) on Illumina NovaSeq S1.

### RNA-seq data analysis

Annotations from Genome assembly GRCm39 (02.09.2020) were adapted by filtering for transcripts with full coverage (defined as having at least one read covering each exon as counted by htseq-count v2.0.4) at least in one of the sample replicates, followed by making intron annotations within the space between two exons of each transcript. Bedtools v2.30.0 subtract and merge were used to first remove sequences with ambiguous (i.e. introns, exons and/or more than one gene) annotations and collapse the remaining annotations to obtain unambiguous annotations for exons and introns for each gene.

Raw RNA-seq reads were adapter- and poly-A trimmed using trim-galore v0.6.10 and poly X and poly G trimmed using fastp v0.23.4, followed by alignment to the reference genome using STAR v2.7.11a. Deduplication and conversion into name-sorted bam files were performed using samtools v1.18. Deeptools bamCoverage v3.5.2 was used with a bin size of 1 and normalization using counts per million to generate stranded coverage tracks. mRNA and pre-mRNA counts were determined as described previously [35], with the following modifications: Htseq-count v2.0.4 was used to count intronic (within intron and intron-exon boundary mapping) and exonic (within exon, exon-exon junction and exon-intron boundary mapping) fragments. Transcripts per million (TPM) values were calculated from raw exonic and intronic counts by normalizing to library size and exon or intron length determined from the collapsed unambiguous genome annotation. Regression analysis was performed comparing the slope and y-intercept for log-log linear models with exonic normalized counts as predictor and intronic normalized counts as response variable each for input and pull-down using the R functions lm, and car::Anova.

TTP target genes were defined as genes with significantly higher counts in pull-down as compared to input. Both intronic and exonic fragment counts were used. Intronic fragment counts were in general low, making them prone to inaccurate determination of significant differences in pull-down as compared to input. To minimize the risk of including potentially false-positives, a threshold for the lowest acceptable counts was employed. The threshold was defined as the average intronic normalized counts in pull-down samples and had the value 3.891038. Genes with normalized intronic or exonic counts (in pull-down) above the threshold were included in the enrichment analysis. Enrichment analysis was performed for intronic and exonic normalized counts separately. Both data sets were first log(ln)-transformed, tested for differences in mean normalized counts of pull-down to input (t-test, two sided) followed by p-value correction using the Benjamini-Hochberg method. Log-fold change was calculated as ln(mean pull-down)-ln(mean input). Significantly enriched genes were defined using the filters padj. ≤ 0.05 and log-fold change > 0. In:Ex ratios are means of the quotients of intronic and exonic normalized counts per sample.

Raw RNA-seq data were deposited at the National Center for Biotechnology Information (NCBI)’s Gene Expression Omnibus (GEO) database under the bioproject identifier PRJNA1031176.

### Expression of recTZF peptides

The TZF domain used for recombinant production in *E. coli* corresponded to a synthetic peptide previously used in binding studies [58]. BL21(DE3) competent cells were transformed with 50 ng of pGEX-4T-1-GST-TEV.TZF-WT or the TZF-NLS^mut^ derivative which were synthesized by Genscript. The constructs contained TTP TZF domain (amino acids 93-165, mouse coordinates) or the mutated TZF version NLS^mut^. Untransformed BL21 was used as empty control. Over night cultures were diluted with fresh LB medium and grown at 37°C until OD_600_ reached ∼0.6. Expression was induced by supplementing cultures with 1 mM IPTG (Sigma), 100 µM ZnSO_4_ and 0.2% glucose (Sigma, Cat # G8270) for 3 h at 37°C at 165 rpm. Afterwards, cells were harvested by centrifugation and lysed by One shot cell disrupter (Constant Systems Ltd.) with a pressure of 2 kbar in lysis buffer (1 M NaCl, 25 mM HEPES pH 7.4, 0.5 mM PMSF, 1x cOmplete protease inhibitor cocktail). Subsequently, lysates were cleared by centrifugation, 6000 rpm 4°C and protein concentration was estimated by BCA protein assay kits (Thermo-Fisher Scientific, Cat #23225).

### RNA Electrophoretic Mobility Shift Assay (RNA EMSA)

RNA EMSA assay was performed using LightShift Chemiluminescent RNA EMSA Kit (Thermo-Fisher Scientific, Cat #20158) according to the manufacturer’s instructions. 5’ biotinylated and unlabeled *Tnf*-ARE probes (5’-AUUAUUUAUUAUUUAUUUAUUAUUUA-3’) were purchased from Microsynth. In brief, HEK293 cells and BL21(DE3) expressing inducible TTP constructs or TZF peptide constructs, respectively, were harvested as previously described. 15 µg of HEK293 cell extract or 100 ng of BL21(DE3) extract were incubated with 0.5 µl Poly-U RNA (100 ng/µl, Sigma, Cat# P9528) 0.25 µl of biotin 5’ labeled *Tnf-*ARE probe (500 nM), 1x REMSA binding buffer, and 5% Glycerol for 25 min at RT. For titration assays, cell lysates were incubated with different ratios of unlabeled to biotinylated probe (0x, 2x, 4x, 6x, 8x, or 10x excess of unlabeled probe). Subsequently, 1X REMSA Loading Buffer was added and samples were run on a 6% non-reducing polyacrylamide gel for 45 min at 100 V in 0.5x TBE. After wet-blotting in 0.5X TBE for 25 min at 250 mA, the Nylon membrane was crosslinked with UV light at 120 mJ/cm^2^ for 60 s using a commercial UV-light crosslinking instrument equipped with 254 nm bulbs. Biotin signal was detected and quantified by chemiluminescence according to standard LightShift Chemiluminescent RNA EMSA Kit protocol using a ChemiDoc™ Touch Imaging System (BioRad).

### TZF structure prediction and modeling

AlphaFold2, USFR ChimeraX and the Robetta server [59–62] were used for structure prediction of TTP TZF defined as region spanning amino acids 93 – 165 (mouse coordinates) corresponding to ZFP36L2 TZF in the PDB entry. ZFP36L2 TZF NMR entry is lacking the most N-terminal and 3 C-terminal amino acids in comparison to the TTP TZF which was kept in the same length as for RNA binding experiments in this study and as used by others previously [58]. Structure prediction using USFR ChimeraX was done without energy minimization. For structure predictions with Robetta comparative modeling PDB entry 1RGO was used as template. Hydrogen bonds were analyzed using UCSF ChimeraX with hydrogen bond parameters set to 0.4 Å distance and 30° angle deviation. Root-mean-square-deviation (RMSD) of the models to 1RGO was analyzed by PYMOLalign with cycles set to 0 for all-atom RMSD calculation. Figures were prepared with PYMOL (http://www.pymol.org/).

### Quantitation and statistical analysis

For all experiments, statistical tests, n, and cut-offs are listed in the figure legend. Statistical analysis was performed using GraphPad Prism v.10.0.2 software.

## ACKNOWLEDGEMENTS

We are grateful to Kevin Eislmayr, Ronja Reinhart, Stefanie Toifl, Josef Laurin and Thomas Peterbauer for fruitful discussions and advice. We thank Florian Ebner and Vitaly Sedlyarov for the generation of TTP-deficient RAW macrophages. All LC-MS/MS measurements were performed on instruments provided by the Vienna Biocenter Core Facilities (VBCF, www.viennabiocenter.org/facilities). The VBCF NGS Facility is acknowledged for RNA sequencing advice.

## EXTENDED DATA FIGURE LEGENDS

**Extended Data Figure 1 (relates to Figure 1).**
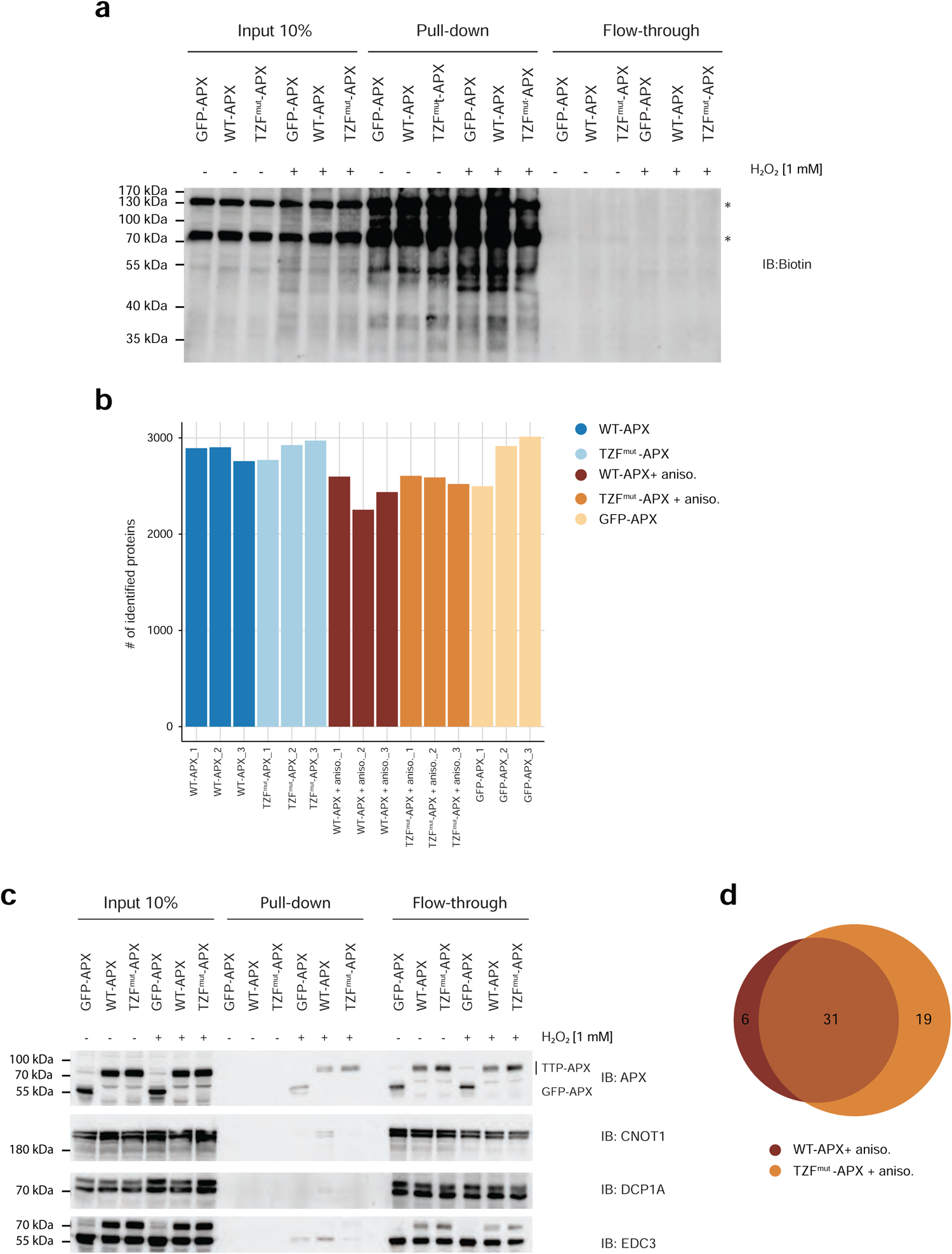
Quality control and validation of proximity labeling. **a.** Biotinylation activity of APX constructs analyzed by Western blotting. WT-APX, TZF^mut^-APX and GFP-APX expression in HEK293 cells was induced (dox, 3.5 h) followed by the addition of biotin-phenol and H_2_O_2_. H_2_O_2_ was omitted, as indicated, to assess endogenous biotinylation. Cell lysates (Input) were subjected to streptavidin pull-down to enrich labeled proteins; pull-down efficiency was estimated by the flow-through analysis. Immunoblotting using biotin antibodies (IB: Biotin) revealed biotinylated proteins. The overall higher biotin signal in H_2_O_2_-treated input and pull-down samples as compared to samples without H_2_O_2_ indicates APX-dependent labeling; biotin signal in samples without H_2_O_2_ treatment indicates endogenous biotinylation; discrete bands (marked with asterisks) present in input and pull-down samples regardless of H_2_O_2_ treatment show positions of the biotin-dependent carboxylases pyruvate carboxylase, mitochondrial (UniProt P11498, upper band) and methylcrotonoyl-CoA carboxylase subunit alpha, mitochondrial (UniProt Q96RQ3, lower band) (**Supplementary Table 1**). **b.** Number of proteins identified in individual replicates by LC-MS/MS. The number of identified proteins in each replicate is calculated from the LFQ intensities before imputation, and reflects peptides mapped to the protein database. **c.** Validation of LC-MS/MS-detected interactions by Western blotting. WT-APX, TZF^mut^-APX and GFP-APX expression in HEK293 cells was induced (dox, 3.5 h) followed by the addition of biotin-phenol and H_2_O_2_. H_2_O_2_ was omitted as indicated to obtain background signals. Cell lysates (Input), streptavidin pull-down and flow-through samples were analyzed by immunoblotting using antibodies to APEX (APX), CNOT1, DCP1A and EDC3. The positions of TTP-APX fusion proteins (WT-APX and TZF^mut^-APX) and the GFP-APX fusion protein are indicated. Note that the signal above the EDC3 band results from incomplete stripping of APX antibodies from the membrane. **d.** Overlap of proteins enriched in WT-APX+aniso (WT-APX+aniso vs. GFP-APX) and TZF^mut^-APX+aniso (TZF^mut^-APX+aniso vs. GFP-APX) interactomes. log_2_ fold change (FC) > 1, padj. ≤ 0.05, n = 3

**Extended Data Figure 2 (relates to Figure 2).**
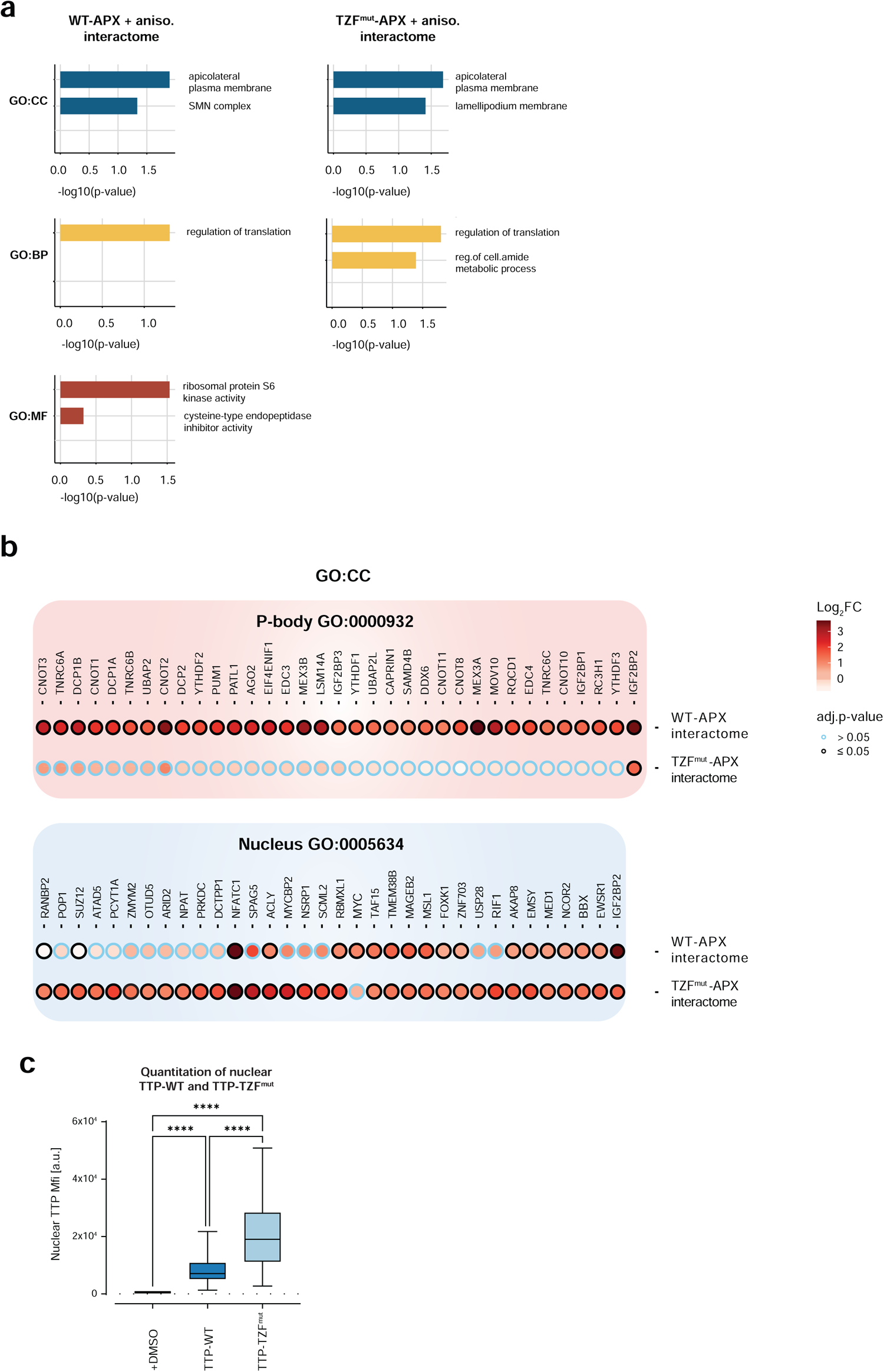
GO term analysis of TTP interactome upon stress, gene-level analysis of representative TTP interactomes, and quantitation of effects of RNA binding on nuclear localization of TTP. **a.** Overrepresentation of GO terms in WT-APX + aniso. and TFZ^mut^-APX + aniso. interactomes. Interactomes were defined as proteins significantly enriched over GFP-APX background labeling control (log_2_FC ≥ 1 and padj. < 0.05, n = 3) in LC-MS/MS data. Bar plots depict significantly overrepresented terms in the GO categories CC, BP and MF. Note that the TZF^mut^-APX + aniso. interactome did not contain any enriched terms in the MF category and is therefore not shown. **b.** Proteins from GO terms P-body (GO:000932) and Nucleus (GO: 0005634) found enriched in WT-APX or TZF^mut^-APX interactomes. Black circles: significantly enriched proteins; blue circles: proteins not enriched. Fold-change (Log_2_FC) enrichment is indicated by color code. **c.** Quantitation of nuclear MFI of TTP-WT and TTP-TZF^mut^. Nuclear masks for quantitation of nuclear MFI were computationally defined by using ImageJ. Box represents interquartile range, horizontal line in box depicts mean. One-way analysis of variance (ANOVA) with Tukey’s multiple comparisons test, n = 62, ****padj. < 0.0001.

**Extended Data Figure 3 (relates to Figure 3).**
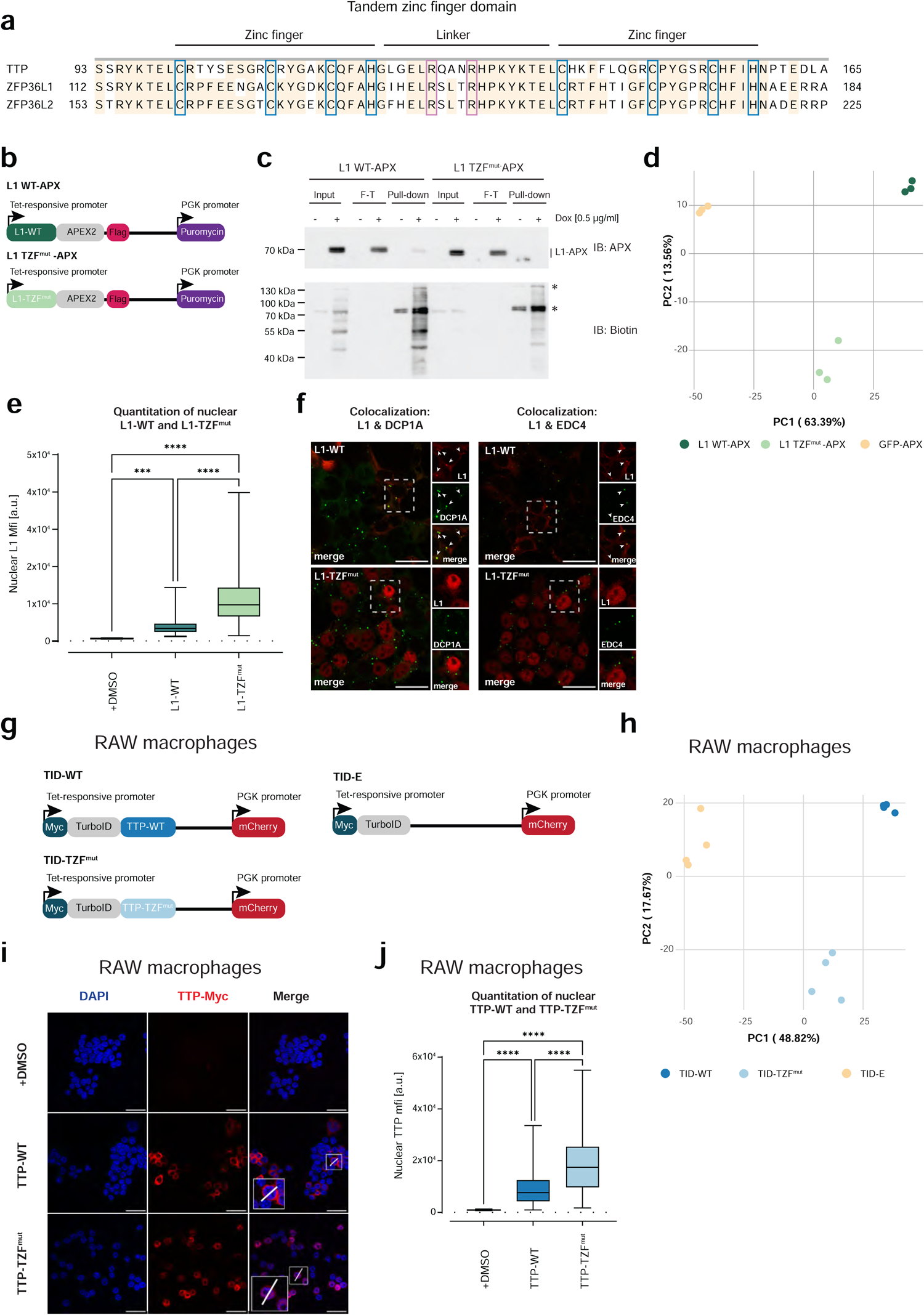
Dependence of interactomes and subcellular localization of ZFP36L1 in HEK293 cells and of TTP in RAW macrophages on RNA binding. **a.** Multiple sequence alignment of TZF domains of murine TTP (NP_035886), ZFP36L1 (NP_031590) and ZFP36L2 (NP_001001806). TZF domain is defined as a 73 amino acid long region comprising zinc fingers (ZFs) 1 and 2, linker between ZFs, conserved N-terminal leader sequence and seven amino C-terminal amino acids, as reported [58]. ZF cysteines and histidines are highlighted in blue; arginines important for the nuclear localization are in lilac. **b.** Schematic representation of ZFP36L1 WT and TZF^mut^ fused N-terminally to APEX2 (L1 WT-APX and L1 TZF^mut^-APX) downstream of a tetracycline-responsive promoter. All constructs contained a C-terminal Flag tag. Puromycin was used for cell selection following lentivirus-mediated transduction of HEK293 cells. **c.** Biotinylation activity of L1 WT-APX and L1 TZF^mut^-APX analyzed by Western blotting. L1 WT-APX and L1 TZF^mut^-APX expression in HEK293 cells was induced (0.5 µg/ml dox, 16 h) followed by the addition of biotin-phenol and H_2_O_2_. Controls without dox treatment were included. Cell lysates (Input) were subjected to streptavidin pull-down (pull-down) to enrich biotinylated proteins; pull-down efficiency was estimated by the flow-through (F-T) analysis. Immunoblotting using biotin antibodies (IB: Biotin) revealed biotinylated proteins. Discrete bands (marked with asterisks) present in input and pull-down samples regardless of H_2_O_2_ treatment show positions of the biotin-dependent carboxylases pyruvate carboxylase, mitochondrial (UniProt P11498, upper band) and methylcrotonoyl-CoA carboxylase subunit alpha, mitochondrial (UniProt Q96RQ3, lower band) (**Supplementary Table 3**). **d.** PCA of protein intensities visualizing clustering of biological triplicates of L1 WT-APX, L1 TZF^mut^-APX and GFP-APX. **e.** Quantitation of nuclear MFI of L1-WT and L1 TZF^mut^. L1-WT and L1-TZF^mut^ expression in HEK293 cells was induced and localization was determined as described in Figure 3c. Nuclear masks for quantitation of nuclear MFI were computationally defined by using ImageJ. Box represents interquartile range, horizontal line in box depicts mean. One-way analysis of variance (ANOVA) with Tukey’s multiple comparisons test, n = 44, ***padj. < 0.001, ****padj. < 0.0001. **f.** Colocalization of L1-WT or L1-TZF^mut^ with DCP1A and EDC4 foci. L1-WT and L1-TZF^mut^ were expressed as in Figure 3c. Immunofluorescence images show L1-WT or L1-TZF^mut^ and endogenous DCP1A and EDC4. Large images depict L1 constructs visualized with a Myc antibody (red); highlighted cells are shown in insets together with DCP1A staining or EDC4 staining (both green) in individual and merged channels. Arrowheads point to colocalized foci of DCP1A or EDC4 with L1. Scale bar = 25 µm. **g.** Schematic representation of TTP WT and TTP TZF^mut^ C-terminally fused to TurboID (TID-WT and TID-TZF^mut^) and allowing doxycycline-inducible expression in RAW macrophages. TID-empty (TID-E) was included as a background labeling control. All constructs were N-terminally Myc tagged. Puromycin resistance served for selection following lentiviral transduction of RAW macrophages. **h.** PCA of protein intensities visualizing clustering of four biological replicates of TID-WT, TID-TZF^mut^ and TID-E. **i, j.** Subcellular localization of TTP-WT and TTP-TZF^mut^ in RAW macrophages. TTP-WT and TTP-TZF^mut^ expression was induced as in Fig. 3h; DMSO was used as vehicle control. (**i**) Immunofluorescence images visualize TTP-WT and TTP-TZF^mut^ detected using a Myc antibody (red); DAPI was used for nuclear staining (blue). Profile lines were drawn through representative cells for MFI quantitation. (**j**) Quantitation of nuclear MFI of TTP-WT and TTP-TZF^mut^ in RAW macrophages. Nuclear masks for quantitation of nuclear MFI were computationally defined by using ImageJ. Box represents the interquartile range, horizontal line depicts the mean. One-way analysis of variance (ANOVA) with Tukey’s multiple comparisons test, n =112, ****padj. < 0.0001.

**Extended Data Figure 4 (relates to Figure 4).**
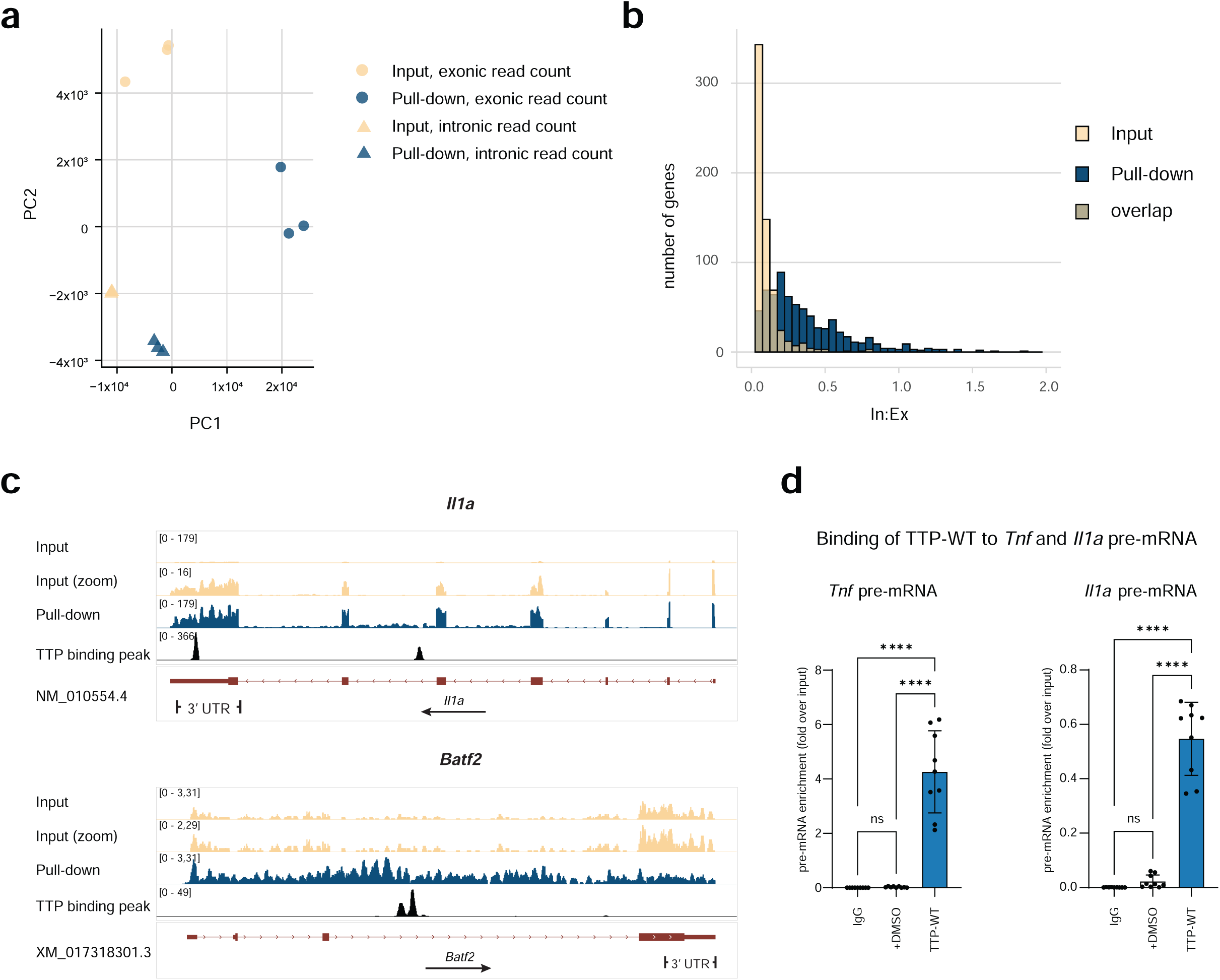
Differential enrichment of pre-mRNA versus mRNA in TTP-bound RNA in RAW macrophages. **a.** Intronic and exonic RNA-seq read counts of TTP-bound RNA (pull-down) and of total input RNA (input) analyzed by PCA. RAW macrophage treatment and RNA isolation were carried out as described in Figure 4b. Following read mapping, exonic and intronic reads were annotated and separately counted. Replicates of individual samples are depicted as indicated. **b.** Distribution of In:Ex ratios across TTP target genes in pull-down and input samples. The histogram visualizes that most genes have an In:Ex ratio of less than 0.1 in input while the pull-down samples show a peak at the In:Ex ratio of 0.2. **c.** Representative IGV tracks for TTP targets *Il1a* and *Batf2* in input and pull-down samples. Data generation and visualization as described in Figure 4. **d.** Analysis of TTP-bound pre-mRNAs by RT-qPCR. Cells were prepared as described in Figure 4b. Cells were treated with 4sU (100 µM) for 7 h to metabolically label cellular RNA, and with LPS (10 ng/ml) for 6 h (i.e. 1 h after 4sU addition) to stimulate expression of TTP targets. Pull-down RNA and input RNA were generated as described in Figure 4b, and subjected to pre-mRNA-specific RT-qPCR analysis. Control analyses were carried out using mouse IgG isotype antibody instead of Myc antibody, and using RAW macrophages treated with vehicle (DMSO) instead of the TTP expression-inducing doxycycline. Pre-mRNA enrichment in TTP-bound RNA shown as fold difference between pull-down and input. One-way analysis of variance (ANOVA), means of RT-qPCR data with SDs, n = 3, each measured in triplicates, ***padj. < 0.001, ****padj. < 0.0001.

**Extended Data Figure 5 (relates to Figure 5).**
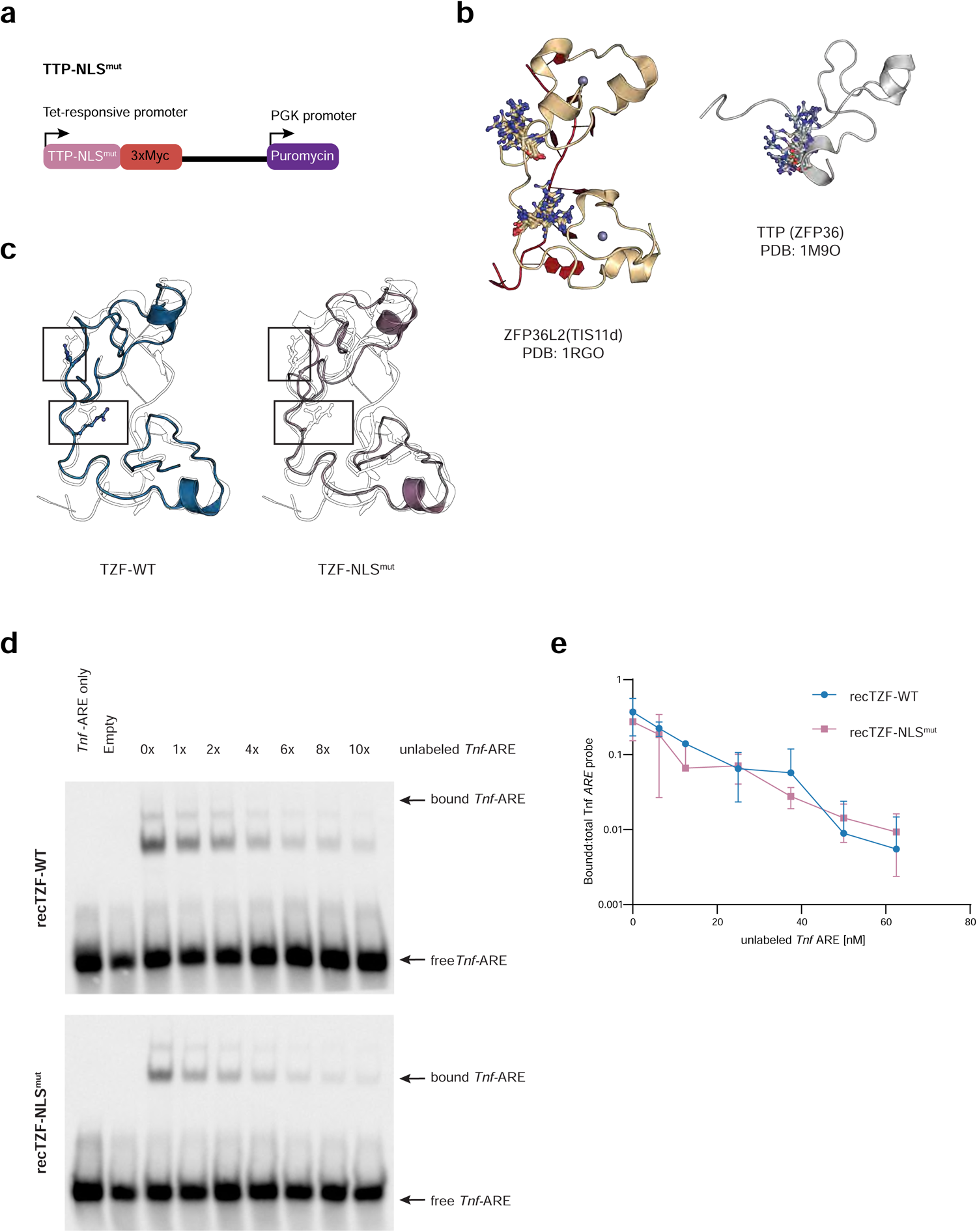
Structural modeling and RNA binding of TZF-WT and TZF-NLS^mut^. **a.** Schematic representation of nuclear import-deficient C-terminally Myc-tagged TTP-NLS^mut^ for doxycycline-inducible expression. Puromycin was used for selection following lentivirus-mediated transduction of HEK293 cells. **b.** NMR ensemble conformations of arginine residues critical for functional NLS in ZFP36L2 and TTP. Reported NMR structure of RNA-bound ZFP36L2 TZF (PDB: 1RGO) and all conformations (20 states in NMR ensemble) of R184 and R188 (full length coordinates) shown on left. Reported NMR structure of TTP ZF1 (PDB: 1M9O) and all conformations (23 states in NMR ensemble) of R126 (full length coordinate) shown on the right. Only R126 is comprised in the available TTP ZF1 structure. Relevant R amino acids shown as balls and sticks, protein backbone structures shown as cartoon. **c.** Superposition of TTP TZF-WT (left, blue) and TTP TZF-NLS^mut^ (right, lilac) AlphaFold2 models with NMR structure of ZFP36L2 TZF (1RGO, state 11, black outline). Protein backbones shown as cartoons, relevant R residues shown as balls & sticks and highlighted in rectangles. Mutations of R to A in TTP TZF-NLS^mut^ (right, lilac) are highlighted in rectangles and visible against R residues in ZFP36L2 background (black outline). **d, e.** Binding of recombinant TZF-WT and TZF-NLS^mut^ GST fusion proteins (recTZF-WT and recTZF-NLS^mut^) to *Tnf*-ARE assessed by RNA EMSA upon different excess of unlabeled *Tnf*-ARE. (**d**) RNA EMSA was performed as described in Figure 5h except for the presence of 0x, 1x, 2x, 4x, 6x, 8x or 10x excess of unlabeled *Tnf*-ARE. Images are representative of three independent experiments. (**e**) Quantitation of RNA EMSA shown in (d). Images were acquired and quantitated using ChemiDoc (BioRad). *Tnf*-ARE-bound fraction was determined as ratio of bound versus total labeled *Tnf*-ARE and depicted as means and SDs (n = 3), in log_10_ scale.

**Extended Data Figure 6 (relates to Figure 6).**
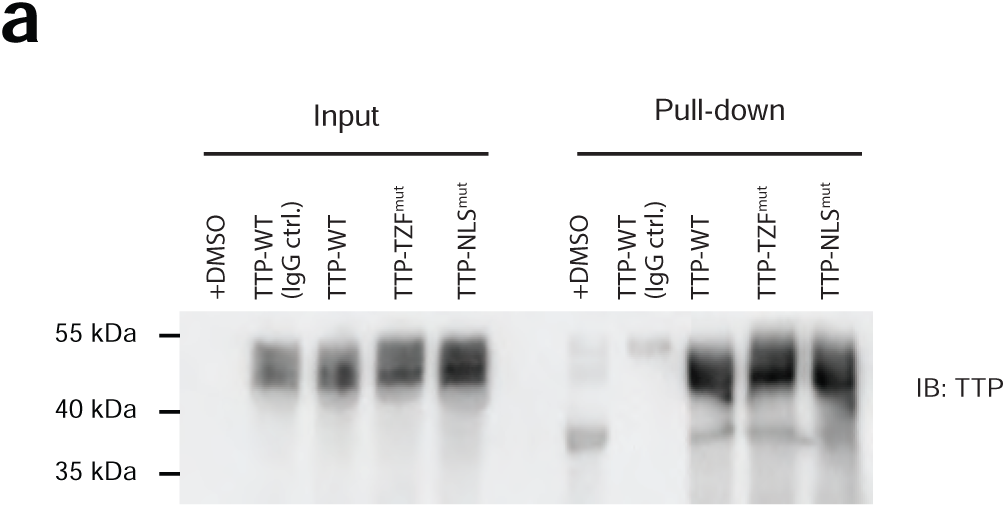
Expression analysis of TTP-WT, TTP-NLS^mut^, and TTP-TZF^mut^ RAW macrophages. Western blot showing protein levels of TTP-WT, TTP-NLS^mut^ and TTP-TZF^mut^ in RAW macrophage cells lysates (Input) and Myc antibody pull-down fractions used Fig. 6a. Control samples include mouse IgG isotype antibody instead of Myc antibody, and using RAW macrophages treated with vehicle (DMSO) instead of the TTP expression-inducing doxycycline. TTP constructs were detected by immunoblotting with TTP antibodies (IB: TTP).

